# Evolutionary expansion of the bacterial *ompW* gene family enables functional diversification during host adaptation

**DOI:** 10.64898/2026.10.09.757860

**Authors:** Alejandro Rubio, Antonio Moreno-Rodríguez, Lucía Ceballos-Romero, Andrés Garzón, Juan A. Marqués-Hinojosa, Elisa Luque-Jiménez, María P. Sánchez-Suero, Jerónimo Pachón, María E. Pachón-Ibáñez, Antonio J. Pérez-Pulido

## Abstract

The outer membrane protein W (OmpW) family comprises minor porins essential for environmental adaptation and host interactions in Gram-negative bacteria. However, *ompW* is conventionally treated as a ubiquitous single-copy gene, masking its true evolutionary complexity. Through comprehensive phylogenetic analysis across thousands of reference genomes, we reveal that *ompW* is present in nearly half of Gram-negative lineages, exhibiting a patchy distribution governed by a birth-and-death model of evolution with recurrent duplications and horizontal transfer. Notably, copy number expansions concentrate in lineages associated with eukaryotic hosts, linking paralog accumulation to niche adaptation. Experimental validation in *Acinetobacter baumannii* reveals four distinct *loci* that maintain high structural conservation despite sequence divergence, yet undergo profound functional specialization spanning from baseline physiological maintenance to modulating host colonization and persistence *in vitro* and *in vivo*. These findings dismantle the single-copy paradigm, demonstrating that widespread gene family expansion and paralog diversification drive the phenotypic plasticity required for bacteria to colonize hostile host environments.

## Introduction

Survival in hostile and changing environments requires Gram-negative bacteria to constantly modulate their outer membrane permeability. The outer membrane protein W (OmpW) family represents a group of minor porins that play a pivotal role in this physiological adaptation, mediating how cells respond to diverse external stress conditions (*1*). Like other outer membrane proteins, its structure is characterized by a beta barrel of eight strands with a hydrophobic inner channel. This channel facilitates diverse metabolic adaptations, including the transport of key nutrient substrates that confer salt tolerance in *Vibrio cholerae* (*2*) as well as growth under anaerobic conditions in *Escherichia coli* (*3*).

While the architectural core of the beta barrel remains highly conserved, the extracellular loop regions exhibit significant sequence variability (*4*). This structural plasticity allows for functional specificity and adaptation to particular ecological and clinical niches, modulating bacterial virulence and host immune evasion (*5–7*).

Beyond its role in baseline physiology and nutrient acquisition, OmpW serves as a primary interface for cellular attacks, acting as a docking site for both antimicrobial agents and bacteriophages. It facilitates the uptake of specific hydrophobic antibiotics (*8*), mediates susceptibility to bacteriocins (*9*), and functions as a receptor for phage infection (*10*). The clinical relevance of this interaction is underscored by an 11 bp deletion in the *ompW* gene consistently found in epidemic *V. cholerae* strains, where the loss of this phage receptor provides resistance against the bacteriophage VP5. Because of this dual nature driving both stress adaptation and vulnerability to external agents, OmpW has emerged as an attractive candidate for vaccine development (*11*). OmpW mediates the attachment of the cystic fibrosis-associated pathogens *Burkholderia cepacia* and *Burkholderia multivorans* to lung epithelial cells, and vaccination with OmpW has been shown to confer protection in mice, preventing infection by these bacteria (*12*). Furthermore, the protein is a promising target for novel antimicrobial strategies, including small molecule inhibitors designed to potentiate existing antibiotics (*13*, *14*).

Among the bacteria where this protein plays a critical role, *Acinetobacter baumannii* stands out as a prominent model for study. This opportunistic pathogen is often multidrug resistant and responsible for numerous hospital outbreaks, making the investigation of OmpW essential for developing new strategies to combat its infections (*15*, *16*). Consequently, this protein has been extensively studied in this bacterium since the early 2000s (*17*–*19*). In *A. baumannii*, OmpW has been shown to be involved in iron uptake and in binding and resistance to antibiotics such as colistin (*8*). Moreover, experiments with *ompW* deletion mutants exhibit diminished virulence traits such as lower adherence and invasion in host cell models (*20*, *21*).

In addition to the multiple phenotypes associated with *ompW* in *A. baumannii*, existing literature presents inconsistent or contradictory findings. While some studies report decreased expression in resistant strains, others observe a significant increase (*8*, *22*, *23*). The presence of previously overlooked multiple copies of the gene within the *A. baumannii* genome could potentially reconcile these discrepancies. Lineage-specific gene duplications may yield distinct gene copies (paralogs) (*24*) that undergo functional divergence, thereby contributing to the observed phenotypic diversity of *ompW*.

To explore the evolutionary history of this gene family, we performed a comprehensive phylogenetic analysis across thousands of bacterial genomes. Our findings reveal that the *ompW* family does not follow a simple vertical lineage, but is instead shaped by a dynamic process of birth and death evolution driven by recurrent gene duplications and horizontal gene transfer. Across diverse lineages, higher copy numbers of *ompW* correlate with host associated lifestyles and pathogenic capabilities. This evolutionary dynamic explains the widespread misannotation in the literature, where *ompW* is conventionally treated as a ubiquitous single copy gene, masking a hidden protein family. Specifically, in *Acinetobacter baumannii*, we identified four distinct paralogs, three of which are present in all genomes, and showed that at least one core gene undergoes horizontal divergence while maintaining high structural conservation. Finally, phenotypic characterization of these paralogs revealed functional diversification, showing contrasting effects on pathogenicity and baseline physiology.

## Results

### Phylogenetic distribution of *ompW* across the bacterial domain

To determine the distribution of *ompW* porin-encoding genes across the bacterial domain, we examined the reference genomes of 16,081 bacterial species, ensuring broad phylogenetic representation. The protein sequences encoded in these genomes were obtained from their corresponding genes, and OmpW proteins were searched for using a reference profile based on 12,482 proteins containing the conserved OmpW domain. As expected, the *ompW* genes were detected exclusively in Gram-negative species (Fig. 1A). They first appear in the families *Chitinophagaceae* and *Sphingobacteriaceae*, with *Filimonas lacunae*, a member of the former family, representing the most basal species known to possess them. In particular, 45.18% of all Gram-negative genomes analyzed (4,173 out of 9,236) had at least one *ompW* gene, whereas only 9.91% (913 genomes) carried more than one *ompW* gene (Fig. 1B). Among these, three bacterial families with 10 or more genomes harboring 4 or more *ompW* genes emerged (Fig. 1C): *Burkholderiaceae* (96 species), *Moraxellaceae* (21 species), and *Microbulbiferaceae* (10 species). Specifically, within these families, the genera *Burkholderia* (38 species), *Paraburkholderia* (34 species), and *Acinetobacter* (21 species) were the most prominent (Fig. 1D).

**Fig. 1.**
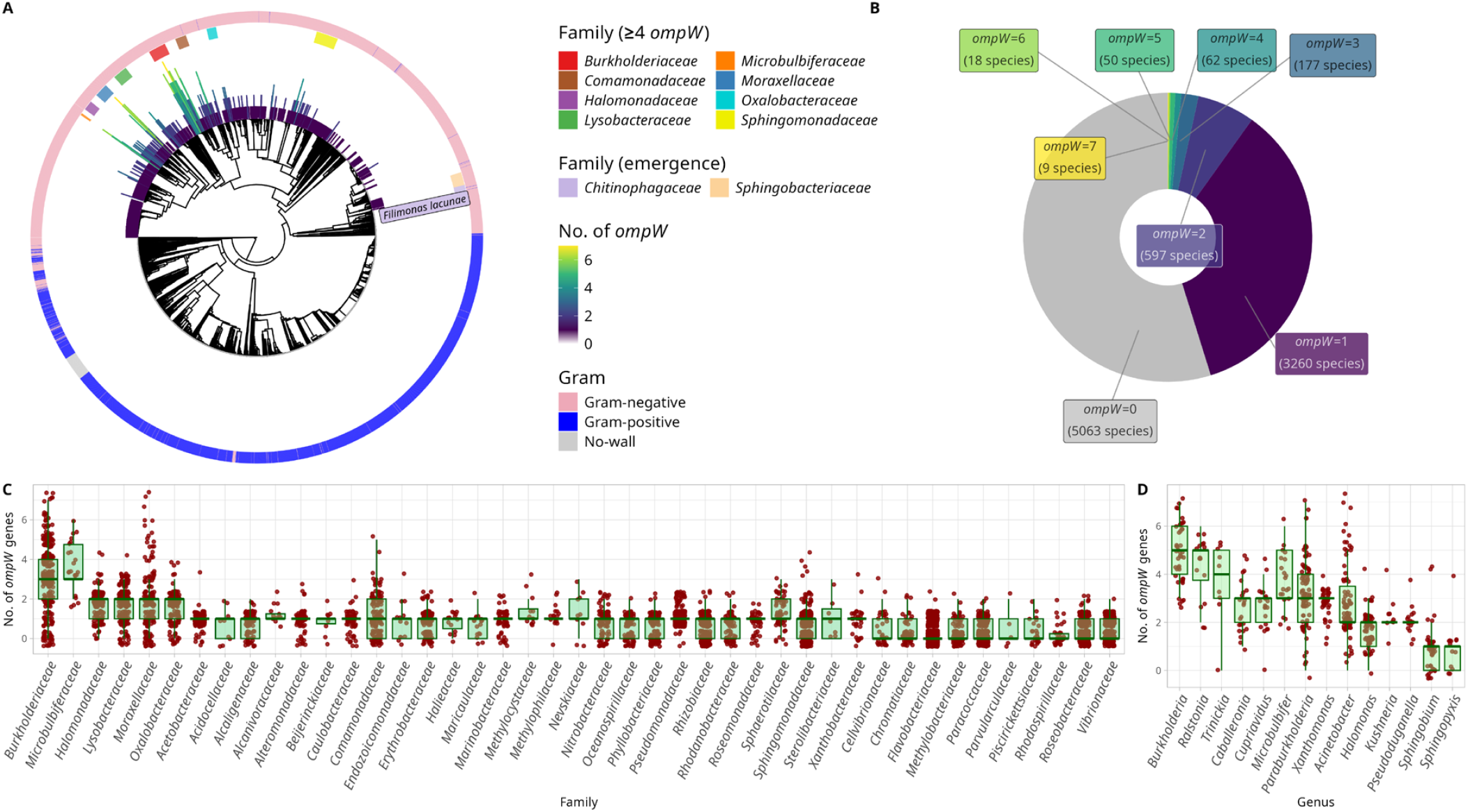
Phylogenetic distribution and copy number of the *ompW* gene in bacteria. (A) Chronogram of bacterial species available in TimeTree (7,783 out of the 16,082 initially considered), highlighting the distinction between Gram-negative and Gram-positive species. The number of *ompW* genes per genome is represented by the bar chart surrounding the tree. Families containing at least one genome with four or more *ompW* copies are highlighted in the inner colored ring, which also marks the two families where the gene likely emerged. The most ancestral sequence was identified in *Filimonas lacunae*, representing the earliest-branching lineage in which the *ompW* gene was detected. (B) Distribution of Gram-negative species genomes according to the number of *ompW* genes they contain. (C) Distribution of the number of *ompW* genes present in the genomes of bacterial families with at least one genome carrying 2 or more genes. (D) Distribution of the number of *ompW* genes present in the genomes of bacterial genera with at least one genome carrying 4 or more genes.

### The *ompW* gene has undergone multiple independent expansions in bacteria

The presence of multiple *ompW* copies across distinct bacterial families could result from either ancestral expansion (monophyletic origin) or independent acquisition and duplication events (polyphyletic origin). To clarify this, we reconstructed a molecular phylogeny using all 5,659 collected OmpW amino acid sequences. The phylogenetic analysis revealed that the ancestral sequence, close to *F. lacunae*, has diversified across the bacterial tree, serving as the ancestral gene for subsequent expansions (Fig. 2A).

**Fig. 2.**
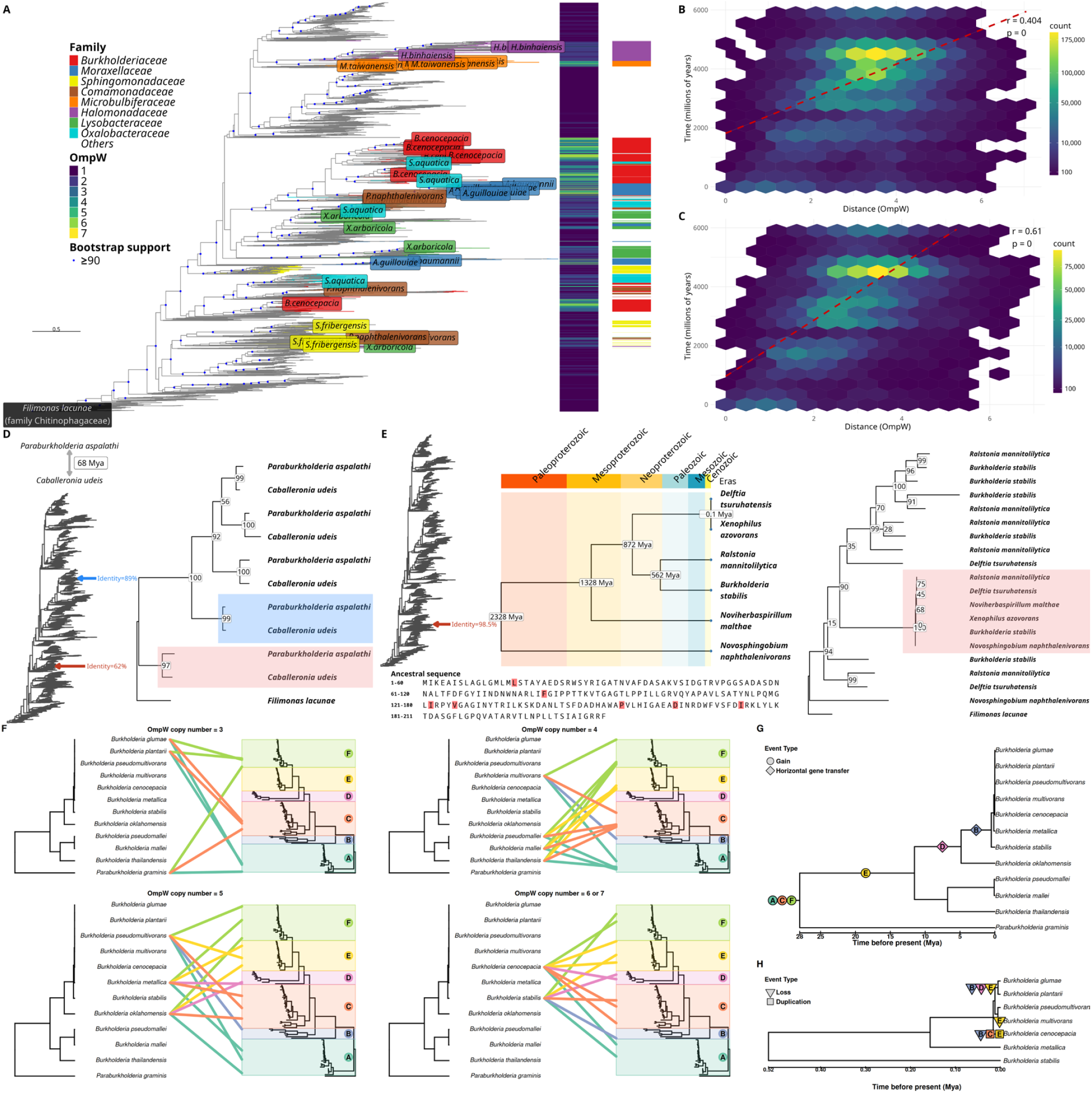
Evolutionary relationships of OmpW proteins and bacterial families with *ompW* gene expansions. (A) Molecular phylogeny of OmpW sequences. OmpW proteins identified in representative species from nine bacterial families containing species with three or more copies of the gene are highlighted. For each family, we selected the species with the highest number of *ompW* copies; *Acinetobacter baumannii* was additionally included. The sequence from *Filimonas lacunae* was used as an outgroup. To the right of the tree, the total number of OmpW proteins per species and their corresponding bacterial family are indicated. (B) Scatter plot of the sequence distance between all pairs of OmpW proteins against the phylogenetic distance between the species that carry them. The points belonging to each pair have been grouped into a density map in which each hexagon represents a count of points. The colors in the legend have been transformed using square root transformation for better visualization. The red line represents the trend line, whose Pearson correlation values and *p-values* are shown in the upper right corner. (C) Same as in (B) but restricted to species harboring a single OmpW copy. (D) Phylogenetic evidence of horizontal gene transfer of *ompW* genes for *Paraburkholderia aspalathi* and *Caballeronia udeis* (diverged ∼68 Mya). The left panel shows a schematic overview of the complete OmpW phylogeny, with colored arrows indicating the relative topological positions of the selected sequences, while the right panel shows the detailed subtree containing the copies of these species. The ancestral canonical copies (red arrow and shading) display a vertical divergence pattern with 62% sequence identity. In contrast, a pair of highly conserved paralogs (blue arrow and shading) exhibit an unexpected 89% sequence identity, strongly suggesting a recent horizontal gene transfer event. (E) Widespread horizontal gene transfer of *ompW* variants across distant evolutionary lineages. The left panel displays a schematic overview of the global OmpW phylogeny, with a red arrow marking a highly conserved cluster sharing 98.5% sequence identity. The central panel shows a chronogram mapping the species divergence times (ranging from 0.1 to 2328 Mya). The right panel shows the detailed subtree with bootstrap support values, highlighting the highly conserved clade shaded in red. Below, the consensus ancestral sequence is shown, with red boxes highlighting the only polymorphic amino acid positions that vary among the six species. (F) Tanglegrams comparing the species chronogram with the OmpW phylogeny for species harboring three, four, five, or six to seven OmpW copies. Colored connecting lines assign each protein to one of the six major OmpW clades (A–F), defined according to the six copies present in *Burkholderia stabilis*. (G) Inferred evolutionary history of these six OmpW lineages across the species chronogram. Colors and letters identify the corresponding OmpW clades from panel F. (H) Enlargement of the recently diverged *B. stabilis* clade, showing lineage-specific losses and putative ancestral duplications that account for the observed variation in *ompW* copy number.

Thus, the phylogenetic analysis uncovered two contrasting patterns of evolution. First, a pattern of vertical inheritance combined with lineage-specific amplifications is observed, in which the ancestral sequence was duplicated, and the resulting copies were transmitted through successive species. This is evident in species from the closely related families *Oxalobacteraceae*, *Comamonadaceae*, and *Burkholderiaceae*. The clustering of several OmpW sequences from these taxa within a single clade supports their common ancestry, likely originating from an ancestral amplification event followed by sequence divergence, which led to the conservation of this gene among closely related lineages. Second, a more recent and localized gene expansion was observed in the later-diverging lineages encompassing the *Microbulbiferaceae* and *Halomonadaceae* families. The emergence of these new gene copies exhibits a synchronized expansion pattern within this specific evolutionary branch rather than independent, isolated events. OmpW sequences from both families cluster tightly together near the apex of the topology, reflecting a shared ancestral expansion or recent divergence that stands in sharp contrast to the more distributed duplications seen in earlier-branching taxa.

In contrast to these massive lineage-specific blocks, other taxonomic groups exhibit a mosaic pattern where ancestral and recently expanded copies coexist within the same family. This coexistence of a vertically retained ancestral copy alongside multiple expanded and highly divergent variants is clearly illustrated by the seven OmpW copies of *B. cenocepacia*. In this species, the ancestral copy clusters with orthologs from closely related species, whereas the remaining copies form a distinct clade that unexpectedly embeds a single copy from an *Oxalobacteraceae* species, a taxon that is phylogenetically distant from *Burkholderiaceae*. On the other hand, within the *Moraxellaceae* family, specifically in *Acinetobacter guillouiae* and *A. baumannii*, one gene copy remains in a branch closer to the root, preserving its ancestral state, while the remaining copies cluster within separate, highly derived branches that show evidence of multiple independent and recent expansions.

Despite these complex dynamics, OmpW sequence divergence retains a discernible baseline signature of vertical inheritance at a global scale. Sequence distances exhibit a moderate positive correlation with the phylogenetic distance between species (Fig. 2B; *r*=0.404, *p-value*=0). This evolutionary signal becomes more pronounced when evaluating exclusively those species that possess a single copy of ompW (Fig. 2C; *r*=0.61, *p-value*=0). These statistical trends suggest that while a primary core copy tends to track vertical bacterial evolution, it coexists with a highly fluid landscape where additional copies frequently arise in specific clades through duplication or horizontal gene transfer, subsequently diverging under alternative selective pressures.

This dynamic pattern is clearly reflected in the frequent occurrence of highly similar sequences in distantly related taxa, whereas markedly divergent sequences can appear within closely related species. Collectively, these findings support a complex evolutionary history shaped by horizontal gene transfer and lineage-specific expansion. To empirically validate these dynamics, we examined specific instances of topological incongruence within the global OmpW phylogeny. This analysis unraveled two distinct scenarios where sequence conservation directly contradicts established host divergence times. In the first case, a localized inspection of *Paraburkholderia aspalathi* and *Caballeronia udeis*, species sharing a common ancestor estimated at ∼68 Mya, revealed a dual evolutionary trajectory for their *ompW* copies. While their ancestral canonical *loci* followed a standard vertical divergence pattern with a baseline 62% sequence identity, a pair of highly conserved paralogs clustered closely together, exhibiting an unexpected 89% sequence identity (Fig. 2D).

This pattern of unexpected sequence preservation across deep evolutionary boundaries was even more pronounced in a second group comprising six highly divergent species (Fig. 2E). The chronogram tracks the divergence times of these hosts back to the Paleoproterozoic era (∼2328 Mya). Despite this substantial temporal distance, these six species harbor an *ompW* cluster that shares a striking 98.5% sequence identity. However, the canonical copy of *ompW* in two of these species, *Xenophilus azovorans* and *Noviherbaspirillum malthae*, has been lost. Taken together, the lack of mutational accumulation in both the *P. aspalathi* and *C. udeis* duplex and the multispecies cluster, combined with the loss of the canonical copy in select lineages, provides robust evidence of dynamic genome remodeling shaped by recent horizontal gene transfers and secondary gene loss.

Birth-and-death dynamics became particularly apparent at finer taxonomic scales. To examine this pattern, we reconstructed the evolutionary history of the *ompW* family within *Burkholderia*, a genus containing some of the highest *ompW* copy numbers in our dataset. The comparison between the species and protein phylogenies revealed substantial variation in copy number and lineage composition, with different species harboring between three and seven copies (Fig. 2F). Our reconstruction suggests that the common ancestor of *Burkholderia* and *Paraburkholderia* already possessed three *ompW* copies. The repertoire subsequently expanded through additional gene gains and horizontal gene transfer events (Fig. 2G). This expansion was followed by lineage-specific losses and duplications, particularly among recently diverged *Burkholderia* species, producing the copy-number variation observed in extant genomes (Fig. 2H).

Extending the analysis beyond *Burkholderia* to the broader *Burkholderiaceae* family revealed a highly discontinuous distribution of the six OmpW lineages, consistent with recurrent gene gains and losses (Suppl. Fig. S1). The most striking pattern involved OmpW lineage B. Within *Ralstonia*, this lineage was detected in only one of the 16 species analyzed, *Ralstonia mannitolilytica*, but its sequence shared 98.6% identity with the corresponding copy from *Burkholderia stabilis*. Such high conservation was unexpected given that *Burkholderia* and *Ralstonia* diverged approximately 562 Mya and therefore strongly supports a relatively recent horizontal gene transfer event, consistent with the phylogenetic evidence presented in Fig. 2E. When the analysis was extended to the complete genus-level chronogram, similarly the conserved B lineage was detected in even more distantly related genera. For example, the protein identified in *Tepidiphilus* shared 98.1% identity with the *B. stabilis* copy, although these lineages diverged approximately 2,328 Mya (Suppl. Fig. S2). Together, these findings suggest that OmpW lineage B has undergone recurrent horizontal transfer across deeply separated bacterial lineages.

Examination of the genomic context provided further support for an ICE mediated transfer mechanism. The B *ompW* gene was located within a putative 116,760 bp integrative and conjugative element (ICE), together with a gene cluster involved in naphthalene degradation and multiple antiphage defense genes (Suppl. Fig. S3). Its association with a mobile element provides a plausible mechanism for the recurrent dissemination of this highly conserved *ompW* variant across deeply divergent bacterial lineages.

### *ompW* genes are more frequent and occur at higher copy numbers in host-associated and pathogenic bacteria

The OmpW protein has been linked to host cell adherence, dissemination, and invasion, which represent key functional traits that underpin the colonization of hostile niches within the host. To test whether an expanded *ompW* repertoire correlates with an increased propensity for host colonization and disease, we classified bacterial species into three distinct categories based on their ecological lifestyle and interaction with the host: pathogens (at least 5 references associating the species with pathogenicity), low-confidence pathogens (1 to 4 references), and non-pathogens (at least 5 references, none of which associated the species with pathogenicity). Analysis of *ompW* copy numbers across these categories revealed that pathogens and low-confidence pathogens typically possess at least one copy (median = 1), whereas non pathogens most frequently lack the gene (median = 0; Fig. 3A). Furthermore, higher copy numbers, particularly those exceeding two, appeared predominantly within species classified as pathogens (Fig. 3B). Consistent with this pattern, species harboring the highest *ompW* copy numbers (5 to 7 copies) were overwhelmingly classified as pathogens or host-associated bacteria, showing a capacity to exploit a diverse range of eukaryotic hosts that includes humans, animals, plants, and invertebrates (Fig. 3C).

**Fig. 3.**
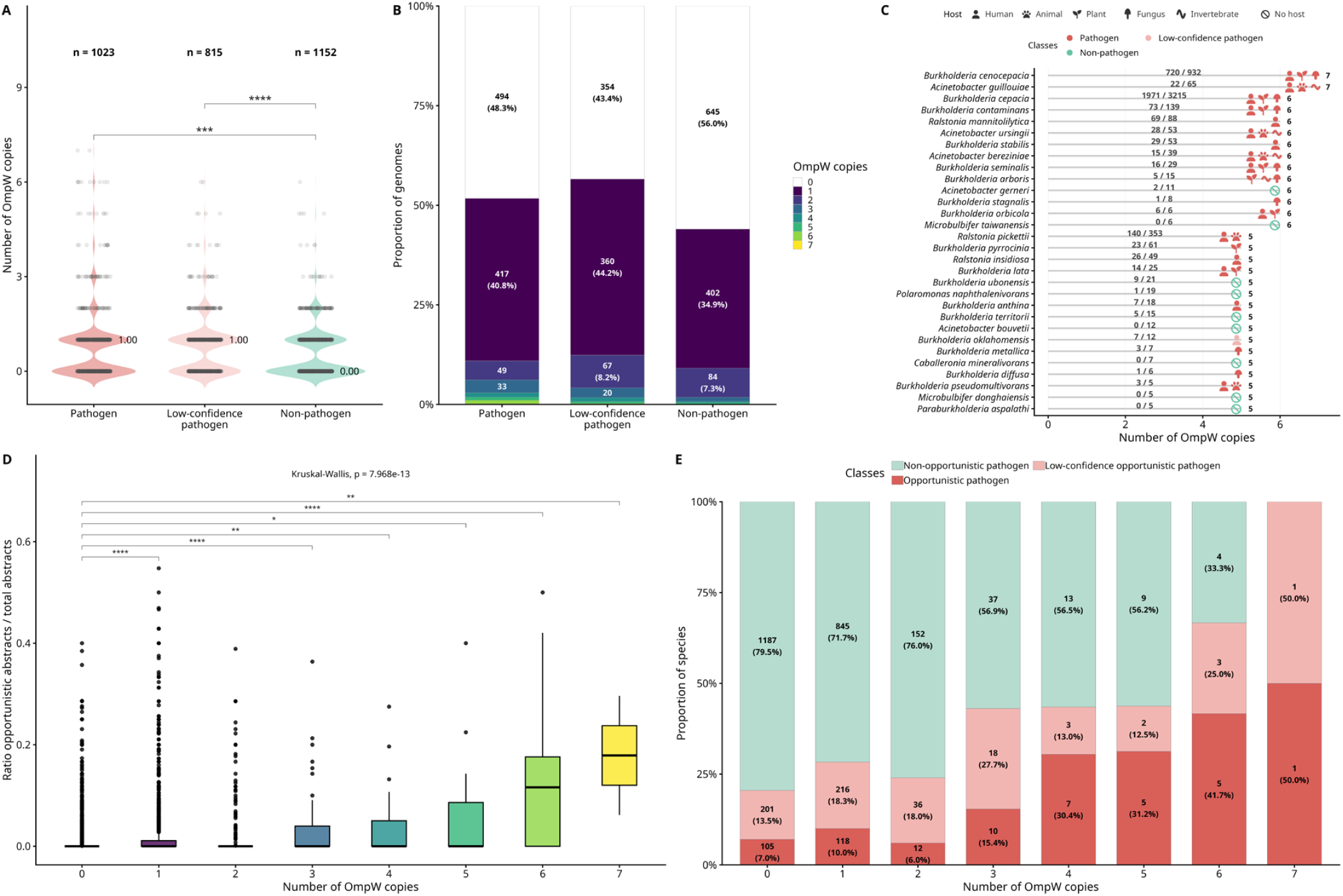
Association between *ompW* copy number and bacterial pathogenicity. (A) Distribution of *ompW* copy numbers among genomes classified as pathogens, low-confidence pathogens, or non-pathogens. Numbers indicate sample sizes and values beside the distributions indicate medians. Significant pairwise differences were assessed using Wilcoxon tests with Benjamini-Hochberg correction. (B) Relative distribution of *ompW* copy numbers within each pathogenicity class. Labels indicate the number and percentage of genomes. (C) Thirty species with the highest *ompW* copy numbers. Colors indicate pathogenicity class, symbols represent host categories manually assigned based on a review of the available literature. These associations include conventional host-pathogen relationships or, in the case of fungi, reported antagonistic or toxic effects exerted by the bacterium on the indicated organism. Values above the lines indicate the number of pathogenicity-related publications relative to the total number of publications retrieved for each species. Numbers on the right indicate *ompW* copy number. (D) Ratio of abstracts mentioning opportunistic pathogenicity to the total number of abstracts for species with different *ompW* copy numbers. The global comparison was performed using a Kruskal-Wallis test, followed by pairwise Wilcoxon tests against species lacking *ompW*, with Benjamini-Hochberg correction. (E) Proportion of species identified as pathogens that can be classified as opportunistic pathogens, low-confidence opportunistic pathogens, or non-opportunistic pathogens according to ompW copy number. Numbers inside the bars indicate species counts and percentages. Asterisks denote adjusted significance levels: *p* < 0.05 (*)*, < 0.01* (**)*, < 0.001* (***), and < 0.0001 (****).

Opportunistic pathogens frequently persist in environmental reservoirs and shift between environmental niches and host associated environments. Such transitions expose bacteria to contrasting conditions and require high ecological flexibility. We examined whether *ompW* copy number was associated with an opportunistic pathogenic lifestyle. Accordingly, *ompW* gene expansion marked a significantly higher propensity toward host opportunism (quantified through relative publication frequency; Kruskal Wallis, *p* = 7.97e-13; Fig. 3D). Consistent with this pattern, opportunistic pathogens constituted a progressively higher proportion of species in expanded copy number categories, showing a marked enrichment in lineages harboring three or more copies compared to single copy or gene lacking genomes (Fig. 3E). These results support a strong evolutionary link between *ompW* gene family expansions and the adoption of an opportunistic pathogenic lifestyle.

### The genomes of *A. baumannii* carry three *ompW* paralogs, including one with two highly divergent variants

One of the bacterial families where *ompW* has undergone the greatest expansion is *Moraxellaceae*. As a prominent member of this family and a clinical model of an opportunistic pathogen capable of surviving in diverse environments, *A. baumannii* represents an ideal candidate to examine this expansion in detail, benefiting from extensive genomic sequence availability driven by intense research interest as well as established tools for genetic manipulation. We therefore analyzed a previously constructed pangenome comprising 7,759 *A. baumannii* genomes.

Although the literature routinely refers to a single *ompW* gene, our pangenome analysis revealed that the *A. baumannii* genome actually harbors up to four distinct *ompW* paralogs located at different chromosomal loci (Fig. 4A). To systematically differentiate these copies throughout this study, we established a numerical nomenclature (*ompW1* through *ompW4*). Three of these paralogs were present as core in virtually all genomes analyzed (*ompW1* = 7,664 genomes; *ompW2* = 7,681; *ompW3* = 7,545), whereas the fourth copy (*ompW4*), belonged to the accessory genome, was found in only a small subset of strains (151 out of 7,759).

**Fig. 4.**
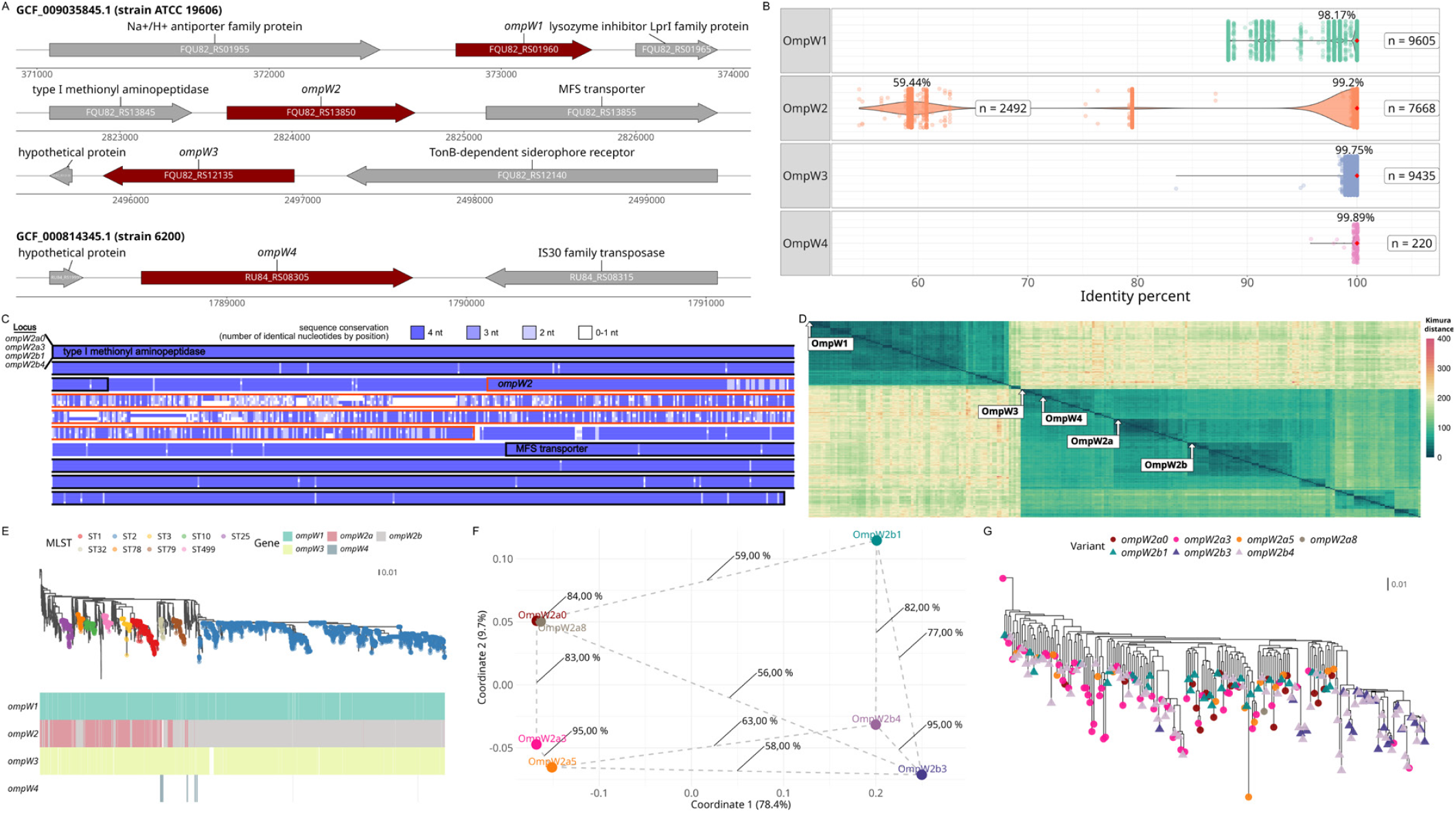
Gene variants of *ompW* in *A. baumannii*. (A) Genomic context of *ompW* isoforms found in *A. baumannii*, illustrating adjacent upstream and downstream flanking genes. (B) Distribution of sequence identity values for all pairs of OmpW proteins in *A. baumannii*, grouped by the four major *loci*. The identity values were calculated using the amino acid sequence of each variant against the reference sequence of its corresponding *locus*. The red dot represents the median. The number of compared sequences and the average identities are shown on each box. In the case of OmpW2, values are separated according to the two distinct groups observed. (C) Sequence conservation of the genomic region encompassing the *ompW2* gene *locus* in *A. baumannii*, including its flanking genes. The alignment incorporates the sequences corresponding to the four most common *ompW2a* and *ompW2b* variants. (D) Kimura distance between different OmpW sequences of the genus *Acinetobacter*. The position of the four variants of *A. baumannii* is marked. (E) Molecular phylogeny showing the presence-absence profile of the different *ompW* genes. The genomes of the most frequent MLST clonal groups are highlighted in the tree. A previous pangenome of the species with 7,759 genomes was used to construct the tree and obtain the OmpW sequences (*64*). (F) Principal coordinates analysis (PCoA) of the four most common OmpW2a and three most common OmpW2b variants based on the amino acid sequence identity matrix. Some selected identity values between variants are indicated as labels. (G) Phylogenetic tree shown in (E), pruned to remove genomes with <0.05 distance and identical *ompW2* variants (7,540 genomes removed). Colored dots show the variant carried in each genome.

Three of these paralogs (OmpW1, OmpW3, and OmpW4) exhibit high sequence conservation. Within each of these paralogs, sequence preservation is remarkably tight: OmpW1 variants display an average identity of 98.17±3.32%, while OmpW3 and OmpW4 remain nearly invariant across strains, with mean identity values of 99.75±0.4% and 99.76±0.36%, respectively (Fig. 4B). In contrast, OmpW2 segregates into two main variant groups (bimodal distribution): OmpW2a, and OmpW2b, which retains approximately 60% mutual identity (global average=89.3±17.7%). This suggests frequent exchange by horizontal gene transfer, which is evident when observing the genomic context of this *locus* (Fig. 4C). While the sequence of the flanking genes of *ompW2* retain 99% identity, the sequence of the four most frequent variants of the *ompW2* gene retain only the sequence encoding the protein signal peptide virtually intact (Fig. 4C; Suppl. Fig. S4-S5).

The *ompW* paralogs in *A. baumannii* appear to have an ancestral origin within the genus, as all four copies are also present in other *Acinetobacter* species, clustering into four distinct group-specific clades (Fig. 4D). However, additional *ompW* paralogs are present in some congeneric species but absent from *A. baumannii*.

Despite the low sequence identity between the OmpW2a and OmpW2b variants, individual genomes harbor either one variant or the other, but never both. This mutually exclusive pattern is expected, as they occupy the same genetic *locus* across *A. baumannii* isolates. Furthermore, while the OmpW2b variant is predominantly found in the high-frequency ST2 clonal group, it is also found interspersed with OmpW2a across other clonal groups (Fig. 4EF). Consequently, even phylogenetically closely related genomes can harbor distinct OmpW2 variants (Fig. 4G), a pattern consistent with frequent replacement by horizontal gene transfer.

### Extracellular hypervariability contrasts with OmpW structural conservation

Because the different OmpW proteins of *A. baumannii* appear to have emerged recently in the *Moraxellaceae* family, high sequence divergence should not be anticipated. Accordingly, OmpW1 remains closer to the ancestral state as the canonical ortholog transmitted via vertical inheritance. This is clearly reflected in amino acid comparisons, where OmpW1 retains 34% identity with the *F. lacunae* ancestor, but shares only 24–28% with the other *A. baumannii* paralogs (Fig. 5A).

**Fig. 5.**
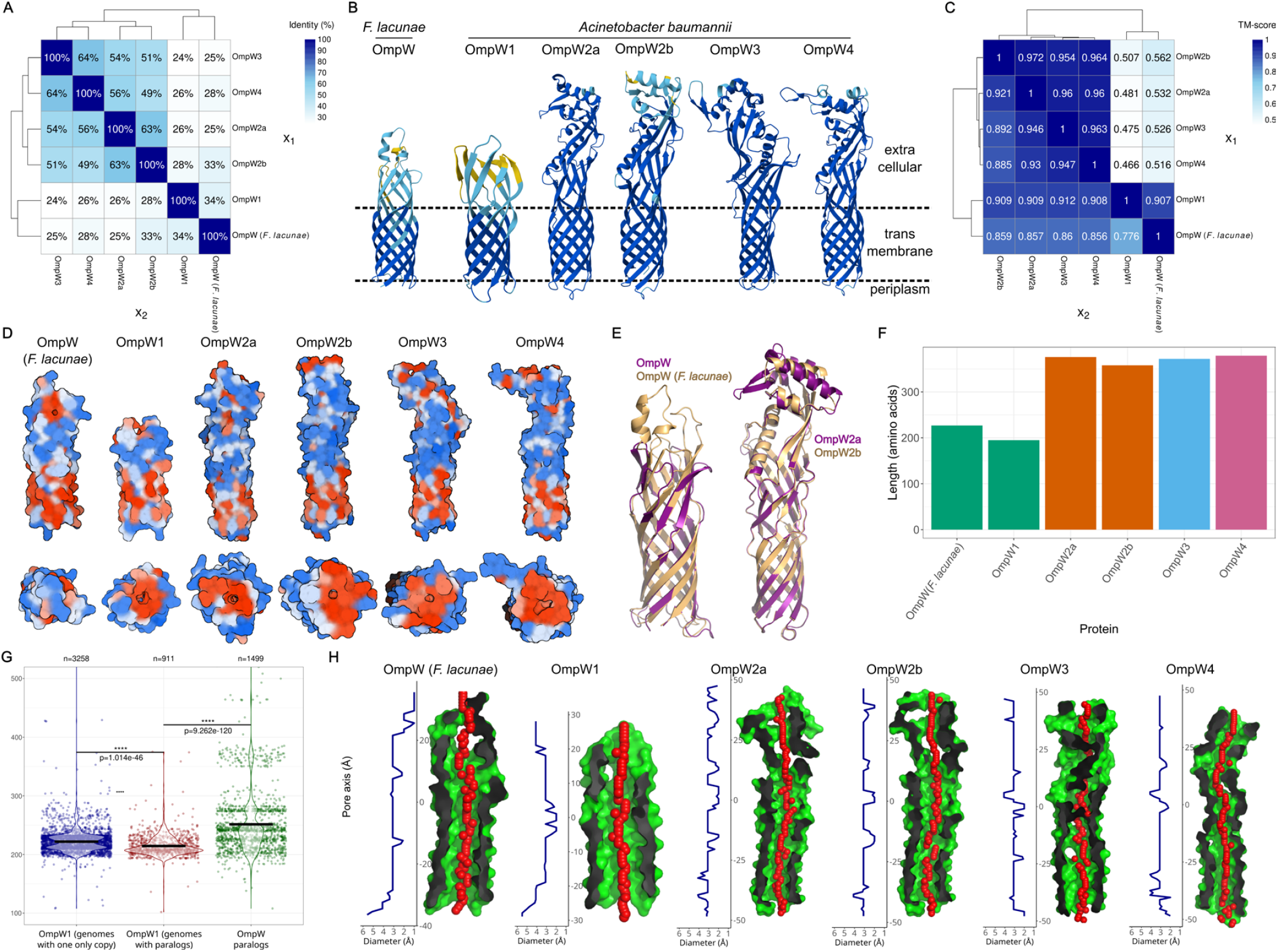
3D structural models of OmpW proteins. (A) Amino acid identity matrix between the different variants of OmpW. (B) 3D structural models of the different variants of OmpW from *A. baumannii* and *Filimonas lacunae*. The colors represent confidence levels predicted by AlphaFold 3: dark blue (very high), light blue (confident), yellow (low), orange (very low). (C) TM-score (Template Modeling score; ranging from 0 to 1) matrix between the different variants of OmpW, computed from pairwise protein structural alignments. Note that the value varies depending on the order of the two sequences (starting with x_1_ or x_2_ may result in different scores). The TM-score is affected by the coverage of the first sequence considered in the structural alignment. (D) Side and top view of the OmpW protein structure models, with hydrophobic regions highlighted in shades of red and hydrophilic regions in blue. (E) Structural alignment of OmpW from *A. baumannii* and *F. lacunae* (TM-score=0.907 and 0.776) and OmpW2a versus OmpW2b from *A. baumannii* (TM-score=0.921 and 0.972). (F) OmpW protein lengths. (G) Distribution of OmpW protein lengths in species with and without paralogs, alongside the length distribution of these paralogous proteins. Significant differences were determined using the Wilcoxon test. (H) Interior space of porins with sequential red spheres marking the internal pathway. Adjacent to each structure is the profile showing the variation in internal diameter along the full length of the porin. The structural representations shown in panel D were generated using BioRender.com.

However, the OmpW2, OmpW3, and OmpW4 paralogs have a greater sequence similarity between them (49-64%), with 63% between the OmpW2a and OmpW2b variants, and 64% between OmpW3 and the rare OmpW4.

Notwithstanding this sequence divergency, the structure of all these proteins is highly similar, especially in their transmembrane region formed by the 8-stranded beta barrel (Fig. 5BC). This is also reflected in the hydrophobicity profile, where highly hydrophobic regions are located not only within the transmembrane domain but also at the extracellular opening of the porin (Fig. 5D). This distribution supports its functional specialization in the transport of hydrophobic molecules. Remarkably, the structure of the two variants of OmpW2, which share only 63% sequence identity, have a virtually identical structural model (TM-score 0.921-0.972 out of a possible maximum of 1) (Fig. 5E).

On the other hand, the structure of OmpW1 is contained within those of the other paralogs and of the *F. lacuae* ancestor (TM-score > 0.9). However, these other proteins are larger due to their more extended extracellular regions, showing only moderate global structural similarity to OmpW1 (TM-scores 0.466-0.507). In other words, the transmembrane regions of all these proteins are structurally nearly identical and differ only in their extracellular domains. These differences are reflected in the protein lengths, with OmpW1 comprising 195 amino acids, while the paralogs range from 358 to 379 amino acids (Fig. 5F).

However, the single OmpW copy harbored by the outgroup species *F. lacunae*spans 227 amino acids in length, featuring a larger extracellular domain than that of *A. baumannii* OmpW1. This observation prompted us to investigate whether OmpW proteins from single-copy species are generally larger to fulfill specialized functions. Conversely, in genomes where paralogs are present, the canonical OmpW1 remains shorter because it strictly retains the core transmembrane porin domain, allowing the additional copies to diversify and adopt functions associated with more expanded extracellular regions. In support of this hypothesis, when comparing the amino acid lengths of OmpW proteins between both groups of species, it was observed that species with a single copy generally have longer proteins than those with multiple copies (222 vs. 214) (Fig. 5G). Indeed, the length distribution of single-copy species displays two distinct modes (218 and 226). This bimodal pattern suggests that a subset of these genomes might actually possess OmpW paralogs that were undetected in the reference strains analyzed in this study, thereby resembling the shorter length typical of multi-copy species, whereas the remaining species genuinely harbor a strict single-copy OmpW. Additionally, the paralogous proteins exhibit greater length variability, ranging from 200 to 500 amino acids, with common clusters around approximate lengths of 200, 230, 240, 280, and 360 amino acids.

Finally, another significant difference between OmpW and its paralogs lies in the inner pore diameter. It is larger in the most ancestral proteins (OmpW1 and OmpW of *F. lacunae*), particularly near the periplasmic region (Fig. 5H).

### *A. baumannii ompW* paralogs display distinct phenotypic roles

To date, a total of 30 published studies have experimentally characterized or investigated *A. baumannii* OmpW proteins. In most of these cases, research has focused specifically on OmpW1, the canonical variant that remains closest to the ancestral state (Fig. 6). However, in at least 7 instances, the term OmpW refers to a paralog encoded by a different *locus*, while in 3 cases, the specific copy remains ambiguous. Among the 7 misidentified instances, only 2 authors explicitly refer to the paralog as *ompW2*. The remaining 5 studies involve genes located at distinct *loci* from the canonical one (OmpW1), but they lack a consistent nomenclature.

**Fig. 6.**
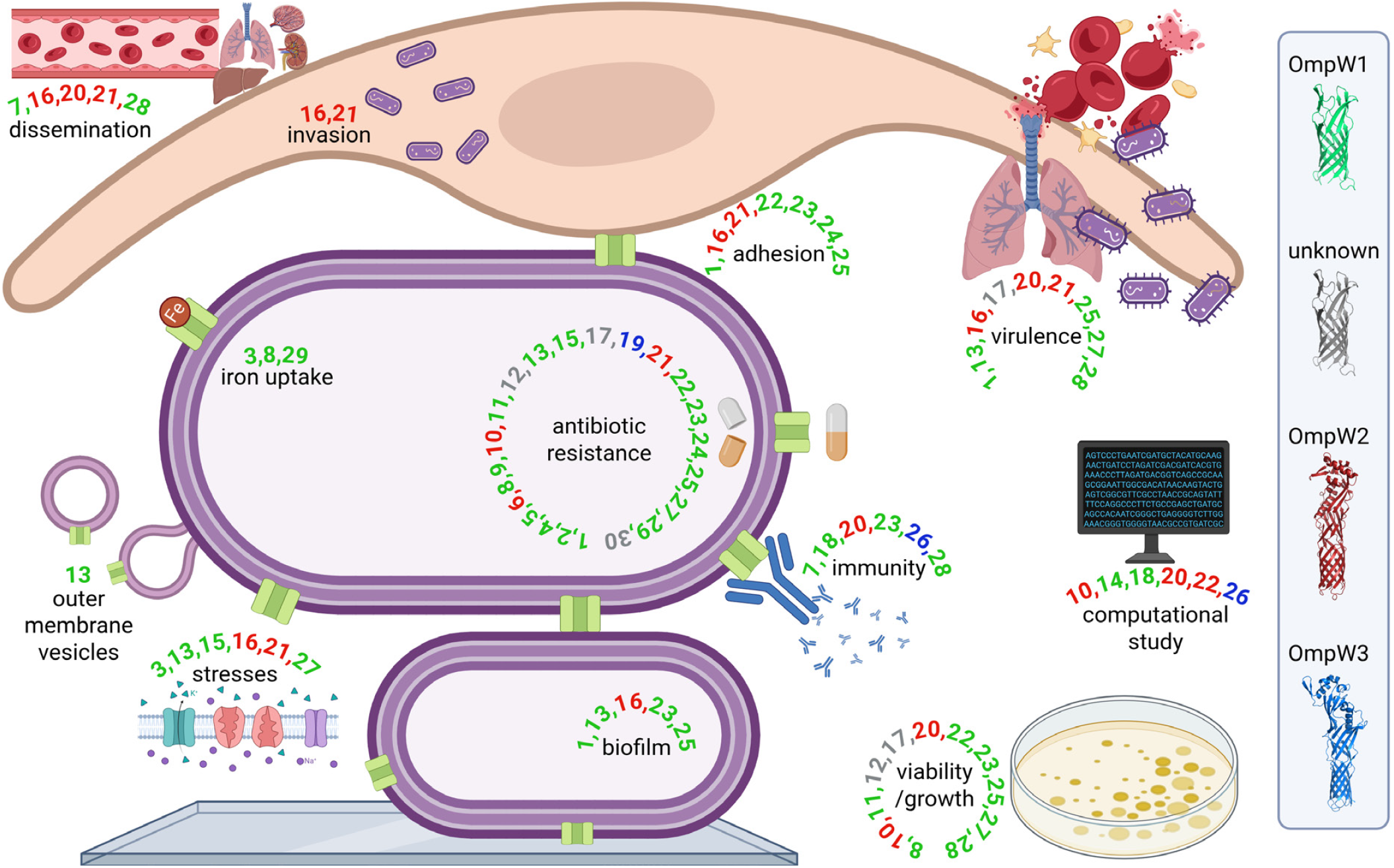
Schematic overview of the phenotypic roles attributed to OmpW variants in the current literature. The numbers associated with each phenotype correspond to specific bibliographic studies, which are detailed in Suppl. Table S1. The color of each number indicates the specific OmpW protein likely investigated in that study, as defined in the legend panel on the right. Figure created with BioRender.com.

In 22 of the 30 analyzed studies, a correlation with antibiotic resistance was reported, identifying it as the most extensively characterized phenotype for *ompW*. Additionally, eleven reports described an impact on bacterial viability or growth, although this effect was likely secondary to antibiotic-induced stress or linked to the iron acquisition response. Furthermore, phenotypes related to virulence are also noteworthy, particularly those involving adhesion and invasion of host cells. Although seven studies implicate *ompW* in adhesion, it is noteworthy that three of the reports focusing on *ompW2* identified phenotypes associated with virulence. Significantly, the invasion phenotype has been exclusively reported in 2 studies that specifically analyzed *ompW2*. Moreover, these three studies on *ompW2* also demonstrated that the protein is implicated in bacterial dissemination to different organs within the infected host.

Finally, studies on *ompW3* remain scarce. To date, this paralog has been investigated strictly in the context of antibiotic resistance and immunity, or has been otherwise limited to *in silico* computational analyses. However, it was mistakenly designated as *ompW* in those studies.

While previous studies have attributed various phenotypes to *ompW*, the specific functional contributions and distinctions among the *A. baumannii* paralogs remain poorly understood. To investigate these phenotypes, mutants targeting the most common variants of the OmpW family were systematically generated using the same parental strain, *A. baumannii* ATCC 17978, specifically disrupting *ompW*, *ompW2a*, and *ompW3* (Fig. 7).

**Fig. 7.**
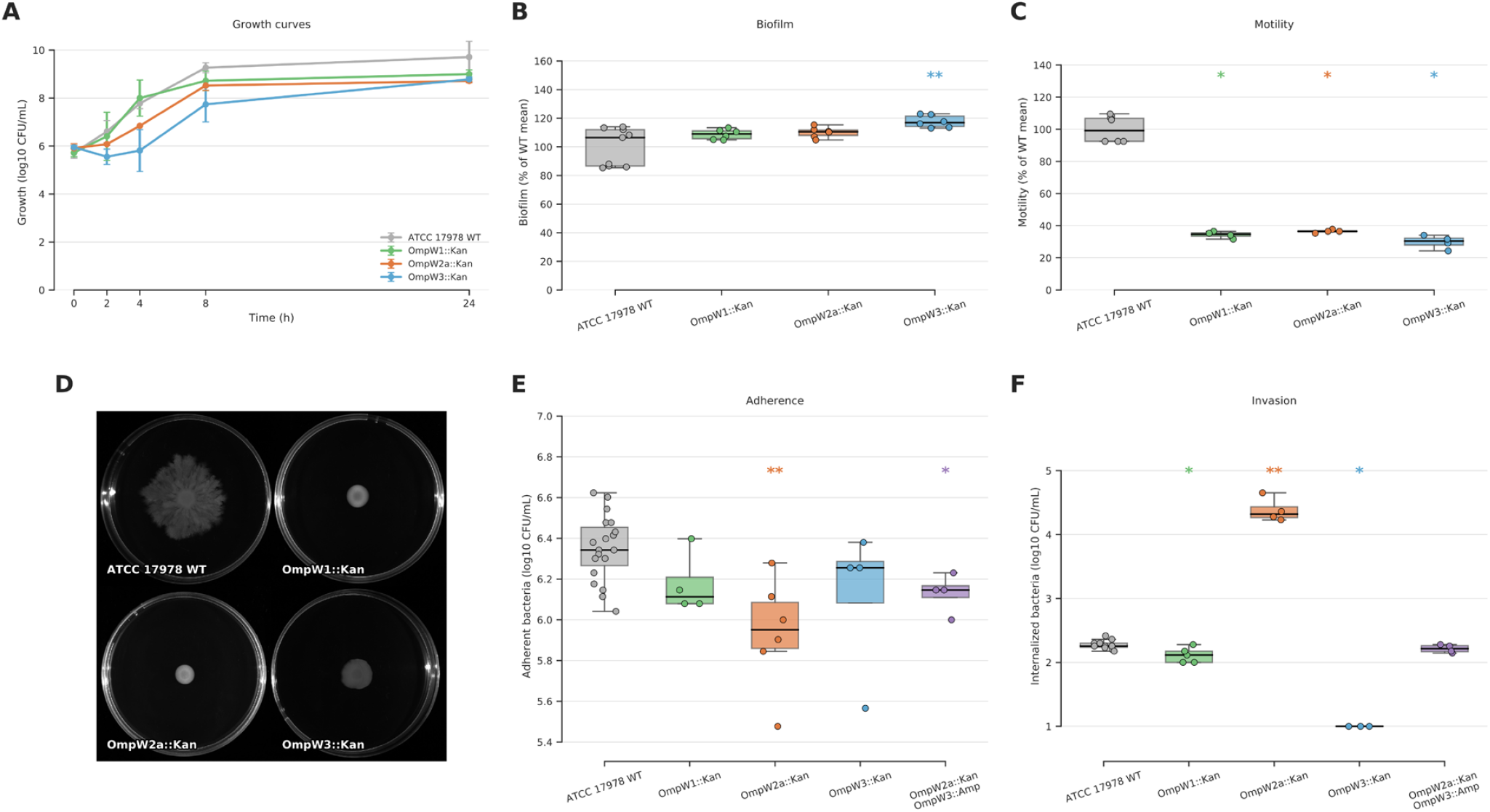
Phenotypic characterization of *A. baumannii ompW* paralog mutants under *in vitro* conditions. (A) *In vitro* bacterial growth assays in Luria-Bertani medium and LB supplemented with kanamycin (50 ng/μL). (B) Biofilm production determined for *A. baumannii* reference strain ATCC 17978 wild-type (WT) as positive control and its isogenic *ompW1*, *ompW2a* and *ompW3* interruption mutants (n=3 in two different days). * p<0.05, Mann-Whitney U test. (C) Effect of *ompW* interruption on surface motility (n=2 in two different days). * p<0.05, Mann-Whitney U test. (D) Motility in *A. baumannii* ATCC 17978 WT, *ompW1*, *ompW2a*, *ompW3*. (E) *A. baumannii* adherence into A549 cells. A549 cells were incubated with *A. baumannii* strain ATCC 17978 WT and its isogenic mutant *ompW1*, *ompW2a* and *ompW3*. Asterisks indicate significant differences (\**p<0.05,* \*\**p*<0.005, \*\*\**p*<0.001) between means with Mann Whitney U test. (F) *A. baumannii* internalization into A549 cells. A549 cells were incubated with *A. baumannii* strain ATCC 17978 WT and its isogenic mutant *ompW1*, *ompW2a* and *ompW3*. Asterisks indicate significant differences (\**p<0.05,* \*\**p*<0.005, \*\*\**p*<0.001) performing Mann Whitney U test.

Our results confirmed that none of the three mutations significantly impacted bacterial growth under standard laboratory conditions. Furthermore, biofilm formation remained largely unaffected across the strains, with the exception of a slight increase observed in the *ompW3* mutant. However, each of the three mutants individually exhibited a significant impairment in surface-associated motility.

On the other hand, we evaluated virulence-associated phenotypes, specifically adhesion to and invasion of human alveolar epithelial cells. Our assays revealed that only the *ompW2* mutant exhibited a significant loss of adherent capacity compared to the wild-type strain. Interestingly, the invasion assays showed divergent results: while internalization was severely impaired in the *ompW3* mutant, the *ompW2* mutant exhibited a dramatic increase in internalization, surpassing the wild-type levels by two orders of magnitude.

A plausible explanation for the internalization result observed in the *ompW2* mutant could be a compensatory mechanism or spatial displacement within the outer membrane. Consequently, the absence of OmpW2 could allow for an increased density or enhanced accessibility of OmpW3, which is the paralog seemingly most implicated in the invasive process, thereby drastically promoting host-cell internalization.

To test the hypothesis that OmpW3 compensates for the absence of OmpW2 and drives the observed hyper-invasive phenotype, we constructed a double-disruption strain (*ompW2*::Kan *ompW3*::Amp) to evaluate both its adhesion and invasion capacities. If OmpW3 is indeed the primary driver of internalization in the *ompW2*-disrupted background, we anticipated that its subsequent inactivation would abrogate the enhanced invasive capacity, bringing it down to levels comparable to those of the single *ompW3*::Amp mutant. Interestingly, rather than completely abolishing internalization, the simultaneous disruption of both paralogs in the *ompW2*::Kan *ompW3*::Amp strain restored invasion strictly to wild-type levels.

*In vivo* infection assays revealed a tissue- and time-dependent effect on bacterial dissemination (Fig. 8). Deletion of *ompW3* produced a global reduction in bacterial burden across all organs analyzed, although this decrease was not uniform. It did cause a significant decrease in peritoneal fluid at 24 hours compared with the wild-type strain. A similar downward trend was also observed for *ompW3* mutant in lung and kidney at 8 hours, although these differences did not reach statistical significance with the sample size analyzed. By contrast, the opposite effect was detected in the other mutant backgrounds, particularly in solid organs such as spleen, lung, kidney and liver, where *ompW1* and *ompW2a* mutants frequently sustained higher bacterial loads than the wild type. This increased accumulation was particularly pronounced in the double mutant strain, which exhibited significantly higher bacterial counts in lungs at 8 hours and in blood at 24 hours. Taken together, these results suggest that the contribution of the different OmpW paralogs to *in vivo* dissemination is not homogeneous, but instead depends on the anatomical compartment and the stage of infection.

**Fig. 8.**
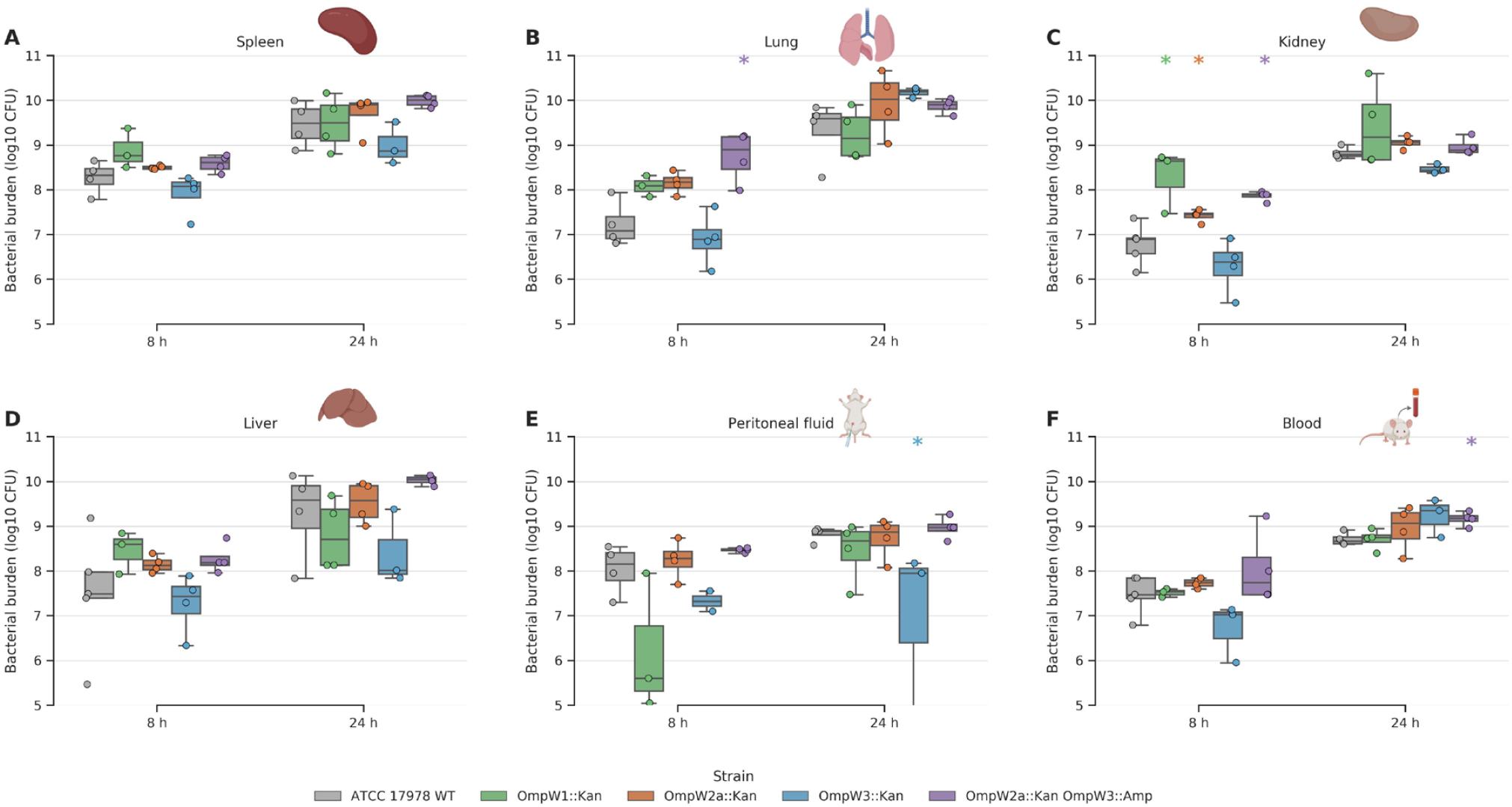
*In vivo* dissemination of wild-type (WT) and *ompW* mutant strains following infection in mice. Bacterial burden was quantified at 8 h and 24 hours post-infection in spleen, lung, kidney, liver, peritoneal fluid, and blood, and is shown as log_10_ CFU. Each point represents one animal. Statistical comparisons were performed against WT at the corresponding time point using a two-sided Mann-Whitney test for organ bacterial burdens. Asterisks indicate statistically significant differences at *p*<0.05. Organ icons were sourced from BioRender.com.

## Discussion

In the present study, a large-scale genomic analysis revealed that *ompW* is present in nearly half of Gram-negative bacterial reference genomes, underscoring its broad evolutionary prevalence across diverse lineages. This widespread yet heterogeneous distribution reflects a global history governed by a birth-and-death model of evolution, where gene duplication and secondary loss operate alongside horizontal gene transfer at distinct taxonomic levels. This complex history explains why sequence divergence retains only a moderate correlation with species phylogeny, as vertically inherited core copies constantly coexist with lineage-specific expansions. However, the true extent of these gene expansions may be underestimated when relying solely on single reference genomes. Notably, while the *A. baumannii* reference genome contains three *ompW* copies, a broader pangenomic analysis of thousands of isolates revealed a fourth paralog that is not universally conserved across all strains, illustrating an accessory nature mediated by recent horizontal gene transfer.

Empirical evidence of these macroevolutionary dynamics is clearly reflected in lineage specific expansions that reshape the *ompW* repertoire, as exemplified by the expansion to up to seven paralogs within the genus *Burkholderia*. The most striking evidence of lateral movement involves an *ompW* lineage within this genus that exhibits up to 98.6% sequence identity across deeply separated genera such as *Ralstonia*, *Novosphingobium* and *Tepidiphilus*, completely contradicting host divergence times that span over 2,300 million years. This extreme preservation across Paleoproterozoic boundaries highlights how birth-and-death evolution operates in tandem with mobile genetic elements, as further evidenced by the location of this gene within a 116 kb integrative and conjugative element carrying naphthalene degradation and antiphage defense systems, a coordinated transport mechanism previously described in the literature (*25*). Crucially, the physical linkage between *ompW* and the naphthalene catabolic gene cluster mirrors genomic configurations observed in polycyclic aromatic hydrocarbon degrading strains such as *Polaromonas naphthalenivorans* CJ2 and *Burkholderia* sp. FM-2 (*26*, *27*). This genomic architecture indicates that *ompW* is co-transferred as a functional component of a larger catabolic operon, thereby imparting complete metabolic and physiological adaptations to recipient lineages in a single lateral transfer event. Beyond a simple mechanism of gene dispersal, this topological configuration illustrates a broader evolutionary paradigm where gene amplification and subfunctionalization allow a protein family to specialize within a specific genetic niche, after which horizontal gene transfer enables distantly related bacteria to rapidly co-opt these pre adapted functional innovations (*28*, *29*). Furthermore, the association of these mobile variants with metabolic and defense traits highlights how recurrent lateral exchange allows opportunistic pathogens to rapidly acquire and preserve fully functional modules across deep taxonomic distances (*30*).

The higher frequency and copy number of *ompW* observed among bacteria capable of causing disease, particularly opportunistic lineages, support an adaptive function for OmpW at the interface between environmental persistence and host colonization. Several experimental studies support this interpretation. In *Acinetobacter baumannii*, *ompW* was strongly upregulated in meropenem-induced persister cells, which also exhibited increased early virulence in a *Galleria mellonella* infection model (*21*). In *Escherichia coli*, OmpW contributes to resistance to phagocytosis and complement-mediated killing, partly through the recruitment of complement factor H (*5*, *31*). OmpW-family proteins have also been implicated in the attachment of members of the *Burkholderia cepacia* complex to lung epithelial cells (*12*), while OmpW contributes to oxidative-stress tolerance in *Actinobacillus pleuropneumoniae* (*32*). Together, these observations suggest that additional *ompW* copies could increase protein dosage or allow regulatory and functional diversification, potentially facilitating survival across environmental and host-associated niches. Nevertheless, our results establish an association rather than a causal relationship, and the concentration of high copy numbers within particular lineages, especially *Burkholderia* and *Acinetobacter*, indicates that phylogenetic history may also contribute to the observed pattern.

OmpW is a porin specialized in the transport of hydrophobic molecules (*1*), which is consistent with our findings demonstrating that the extracellular exit channel across all *A. baumannii* paralogs is predominantly hydrophobic. Here we have found that its eight-stranded beta-barrel is also conserved in all paralogs, despite their great divergence at the amino acid sequence level. However, the extracellular region differs among the various paralogs, suggesting their specialization for different functions, especially in this region that is most in contact with the environment. This specialization is particularly evident in the OmpW paralogs, as their extracellular region is extended by about 200 amino acids compared to the canonical OmpW protein. This conservation has already been demonstrated in the case of the OmpW protein structures of *E. coli* and *Klebsiella pneumoniae* (*4*). However, the differences in their extracellular region do not prevent OmpW from acting in the reception of plasmids between both species.

In *A. baumannii*, the canonical OmpW1 protein, alongside OmpW3 and OmpW4, exhibits high sequence conservation across strains. In contrast, OmpW2 exists as two highly divergent variants distributed throughout the species. While these two internal OmpW2 variants share only 60% amino acid identity with each other, they are predicted to maintain an almost identical structural fold. This divergence in primary sequence, coupled with structural stability, suggests a specialized evolutionary path for the OmpW2 lineage. At the gene level, the region corresponding to the signal peptide is also almost identical, pointing to a common export pathway to the outer membrane, with the rest of the gene being highly divergent at the nucleotide sequence level. This fact could be explained by the mosaic evolution theory of bacterial outer membrane proteins, which posits that proteins evolve as combinations of independent structural modules subjected to different evolutionary pressures (*33*). This modular dynamic is well documented in *E. coli* outer membrane proteins, where variable extracellular loop regions preserve overall structure and function while enabling adaptation through recombination and positive selection against phage predation or immune resistance (*34*).

This OmpW2 variant has been previously analyzed in the literature, and it was correctly designated as a distinct protein from the canonical OmpW in at least two studies (*13*, *35*), whereas in other instances it was erroneously identified as OmpW itself (*20*, *36*, *37*). This lack of resolution is further compounded by the micro-heterogeneity of the protein family, as early proteomic characterizations of *A. baumannii* cell envelopes identified up to four distinct spots for OmpW attributed to post-translational modifications (*17*). The presence of multiple variants of the canonical protein, alongside the co-existence of highly divergent genomic paralogs, underscores the extreme complexity of this outer membrane family and reinforces the need for precise sequence validation.

Multiple studies with OmpW have demonstrated its relevance in virulence and antibiotic resistance (*8*, *13*, *17*–*19*, *21*–*23*, *3C*–*4S*), as well as its involvement in other biological processes such as biofilm formation (*17*, *20*, *22*, *37*, *47*), adhesion (*17*, *20*, *23*, *37*, *44*–*46*) and invasion of eukaryotic cells (*20*, *37*), and the uptake of multiple compounds such as iron (*8*, *48*, *50*). In line with its role in drug susceptibility, recent findings in carbapenem-resistant *A. baumannii* isolates indicate that the upregulation of outer membrane porins, including *ompW*, increases membrane permeability and facilitates intracellular antibiotic accumulation, thereby enhancing meropenem lethality (*47*). However, here we have shown that phenotypes are sometimes described without certainty as to which paralogous gene is being referred to. This omission is particularly evident in studies evaluating broad outer membrane responses to antimicrobial treatments, where *ompW* expression is often discussed as a single entity despite the presence of multiple highly divergent paralogs in the genome. This highlights the importance of accompanying gene and protein names with unique identifiers that allow us to know with certainty which sequence has been used.

In parallel, the significant loss of adherence observed in our *ompW2* mutant highlights the critical, yet complex, role of this specific paralog in host-cell interactions (*20*, *37*). The functional relevance of the OmpW family in mediating bacterial attachment to eukaryotic surfaces is well documented in other Gram-negative species, as demonstrated by the single-copy OmpW that contributes to host cell adherence and colonization in avian pathogenic *E. coli* strains (*51*). In *A. baumannii*, this involvement appears to be highly dynamic. A recent preprint examining longitudinal expression data from the transcriptomic interplay between *A. baumannii*, human macrophages, and polymyxin reveals a dynamic regulation pattern for *ompW2*, characterized by an initial upregulation that subsequently shifts to a marked decrease by 24 hours post-infection (*37*). This temporal shift suggests that rather than acting as a static structural adhesin, OmpW2 functions within a tightly coordinated window during the early phases of colonization. Intriguingly, this matches transcriptional models such as the mentioned pathogenic *E. coli*, where *ompW* is actively modulated by oxygen-responsive regulators like EtrA in response to environmental stressors like hypoxia, further explaining why its absence permanently impairs the stabilization of host-cell attachment over time in our end-point assays.

Regarding the double-mutant responses, the simultaneous deletion of both paralogs restored invasion to wild-type levels. This unexpected phenotype suggests a complex biophysical compensation mechanism within the bacterial outer membrane. Given that these porins occupy significant physical space, their concurrent absence likely triggers a major structural reorganization or alterations in membrane fluidity. Consequently, this membrane destabilization could enhance the exposure or accessibility of alternative surface factors, such as the canonical OmpW, thereby restoring basal host-cell invasion levels through an uncharacterized compensatory pathway. Such structural and biophysical reorganizations within the cell envelope match observations in other outer membrane protein families where porin density directly modulates lipid-protein packing and surface component accessibility (*52*, *53*).

Apart from facilitating initial colonization, OmpW porins are also essential for surviving subsequent host defenses. In *E. coli*, the canonical protein confers resistance to innate immunity by reducing phagocytosis (*5*). In the case of *A. baumannii*, our *in vivo* infection assays demonstrated that the deletion of *ompW3* led to a global reduction in bacterial burden in the analyzed organs. This suggests that following gene expansion events within this species, OmpW3 may have specifically evolved or maintained this protective role, acting as a determinant factor for evading host clearance during systemic dissemination.

A striking finding in our study was that the individual deletion of *ompW1*, *ompW2*, or *ompW3* resulted in a consistent loss of surface-associated motility. Given that these paralogs have significantly diverged in sequence, this shared phenotype suggests a functional interdependence rather than redundancy. It is possible that each OmpW variant plays a distinct but essential role in maintaining the specific physicochemical properties and integrity of the outer membrane required for surface translocation. This biophysical requirement is strongly supported by recent comprehensive evaluations of OmpA-like porins, such as OmpA, ArfA, YiaD, and CarO, in the *A. baumannii* AB5075 strain, where the individual deletion of each porin consistently abolished or heavily reduced surface motility while exerting a much more restricted, gene-specific impact on biofilm formation (*52*). Alternatively, these proteins might be independently required for the correct assembly or coordination of Type IV pili. Since surface motility in this species relies on mechanical pilus retraction governed by genes like *pilT* (*54*), the absence of any single OmpW component might cause a localized membrane destabilization that disrupts the precise mechanical extension or retraction processes necessary for movement.

In conclusion, our study underscores a broader evolutionary strategy in Gram-negative bacteria, where outer membrane porin expansion drives functional specialization for both environmental adaptation and host interaction. While certain paralogs modulate surface translocation and initial attachment, others evolve to ensure survival during dissemination. Ultimately, these overlapping yet distinct roles highlight the urgent need for robust gene notation standards to accurately identify specific sequences, a critical step for fully understanding outer membrane biology across diverse ecosystems.

## Materials and Methods

### Bacterial genomes and identification of OmpW sequences

The 16,081 reference genomes of bacteria from the NCBI Genome with date March 2025 were downloaded (Suppl. Table S2). The family and order to which the species belonged was obtained from NCBI Taxonomy, and the most probable Gram stain classification was obtained through BacDive 2025 (*55*) or by specific bibliographic searches of the family, genus, or species. TimeTree 5 was used for species phylogeny (*56*), finding divergence times for 7,868 species. After removing ambiguous species labeled *Candidatus* and *Incertae Sedis*, a total of 7,783 genomes were finally retained in the phylogeny.

Structural annotation of all starting genomes was performed using Bakta v1.10.3 (*57*) to predict protein-coding genes. To identify OmpW proteins, a hidden Markov model profile was constructed with HMMER v3.4 (*58*) utilizing all sequences associated with the IPR005618 domain from the InterPro database (*59*) (Suppl. Fig. S6). The domain amino acid sequences were aligned using MAFFT v7.526, and the resulting profile was searched against the predicted proteomes. An E-value threshold of 1e-30 was selected to filter hits; this restrictive threshold was implemented to mitigate false positives while ensuring an exhaustive retrieval of all true OmpW variants.

### Molecular phylogeny of OmpW proteins

The amino acid sequences of all OmpW proteins found in the bacterial genomes were taken, and a multiple alignment was performed using MAFFT v.7.526 and the FFT-NS-1 model (*60*). The alignment was further refined by removing regions with excessive gaps, few informative sites, and duplicate sequences using ClipKIT v2.2.4 with the kpi-smart-gap method (*61*). Then, the phylogeny was built using the IQ-TREE v2.4.0 algorithm, applying the Q.pfam+F+G4 model and 1,000 bootstrap replicates(*62*). The optimal model was determined with the integrated ModelFinder algorithm in IQ-TREE.

Phylogenetic distances between sequences were calculated using the cophenetic function from the R library stats (v4.5.2). These values were used as a proxy for protein divergence across the OmpW family. To distinguish between primary orthologs and paralogs, the sequence displaying the shortest distance to the *F. lacunae* ortholog was designated as the ancestral-like OmpW, while any additional copies were classified as OmpW paralogs. In parallel, divergence times between species (expressed in millions of years) were retrieved from the TimeTree 5 database. This allowed for a direct comparison between species-level evolutionary history and the sequence divergence rates of the OmpW family.

### OmpW evolutionary dynamics and mobile element analysis

To investigate *ompW* gain-and-loss dynamics, the OmpW protein phylogeny was compared with TimeTree-derived species chronograms using ape and phytools in R.

Tanglegrams linked each OmpW sequence to its corresponding species and grouped taxa according to copy number. Gene gains, losses, duplications and putative horizontal transfer events were inferred from incongruences between the protein and species trees, sequence conservation and the distribution of the OmpW lineages.

For the supplementary analyses, orthologs of the six *Burkholderia stabilis* OmpW proteins were identified by reciprocal BLASTP. Hits were retained when identity was ≥20% and subject coverage was ≥85% in both search directions. Mean amino acid identity and the proportion of species carrying each ortholog were calculated for every genus and displayed alongside genus level chronograms.

The genomic region carrying the horizontally acquired B *ompW* copy was analyzed using ICEfinder 2.0 through the ICEberg 3.0 web server (*63*). Additional Vir and conjugation-associated proteins were identified by BLASTP analysis of surrounding open reading frames. The element was provisionally classified as a putative ICE based on its predicted attachment sites, integrase, relaxase and conjugation-related genes (Suppl. Fig. S3).

### Pathogenicity, opportunism and host association

Associations between *ompW* copy number and bacterial pathogenicity were evaluated using literature information retrieved from PubMed through the NCBI Entrez API (accessed July 2026). Searches were performed using each species name in the title or abstract fields. The total number of publications was recorded, together with the number of records containing predefined terms related to pathogenicity (including *pathogen*, *disease*, *infection* and *virulence*) or opportunistic infection (including *opportunistic*, *immunocompromised host* and *underlying disease*). Only species represented by at least five PubMed records were retained.

Organism associations and pathogenic lifestyles were manually assigned for the 30 bacterial species displaying the highest *ompW* copy numbers through a comprehensive and manual review of the available literature. Species were categorized by host/organism association as human, animal, plant, fungus, or invertebrate. These associations encompassed conventional infection or colonization relationships as well as experimentally reported antagonistic or toxic effects exerted by the bacterium against the indicated organism (particularly for fungi and selected invertebrates). Additionally, species were classified into pathogenic, opportunistic pathogenic, or non-pathogenic lifestyles based on established clinical and ecological evidence. Species lacking documented evidence for specific interactions or virulence were classified as having no reported association.

Organism associations were manually assigned through a review of the available literature and categorized as human, animal, plant, fungus or invertebrate. These associations included conventional infection or colonization relationships and, for fungi and selected other organisms, experimentally reported antagonistic or toxic effects exerted by the bacterium on the indicated organism. Species for which no such evidence was found were classified as having no reported association.

### Pangenome of Acinetobacter baumannii

To evaluate the distribution of *ompW* paralogs at the species level, we utilized a previously described pangenome of *Acinetobacter baumannii* comprising 8,929 genomes (*64*). These isolates were categorized according to multilocus sequence typing (MLST) following the Pasteur scheme. To identify *ompW* orthologs and paralogs, we conducted a profile-based search using HMMER v3.4, applying the specific hidden Markov model (HMM) profiles and significance thresholds previously established to the protein sequences. To categorize the resulting protein sequences, we performed clustering analysis using CD-HIT v.4.8.1 (*65*). Sequences were grouped into distinct variants based on a minimum identity threshold of 90%, ensuring a robust classification of the protein family. Each cluster generated by CD-HIT was defined as a distinct protein variant. To further investigate the sequence space and diversity among OmpW2 variants, we performed a Classical Multidimensional Scaling (MDS) analysis, utilizing the cmdscale function in the R programming language. The genome of *A. baumannii* ATCC 19606 (NCBI Accession: GCF_009035845.1) was employed as the reference for genomic comparisons and variant positioning.

To construct the molecular phylogeny, we selected the amino acid sequences of genes present in 99% of the analyzed genomes. This resulted in a total of 589 core proteins. The sequences were concatenated by strain and aligned with MAFFT v7.271 using the E-INS-i option (*60*). The alignment was further refined by removing regions with excessive gaps, few informative sites, and duplicate sequences using ClipKIT v2.2.4 with the KPI method (*61*). The molecular phylogeny was then inferred with IQ-TREE v2.3.4, applying the VT+R10 model and 1,000 bootstrap replicates (*62*). The optimal model was determined with the integrated ModelFinder algorithm in IQ-TREE. The tree was visualized with the R package ggtree v1.10.5 (*CC*). The average branch length obtained was 0.0034 ± 0.0134 across a total of 13,227 branches (maximum length: 0.9356, minimum length: 0.000001). This corresponds to an average of 1.34 substitutions per position across the 589 proteins analyzed. The tree was pruned by comparing all pairs of contiguous genomes and removing redundant genomes that were separated by a minimum length of 0.05 and had the same OmpW2 variant.

To create the identity matrix, all previously identified OmpW sequences belonging to the *Acinetobacter* genus were aligned using the MAFFT algorithm FFT-NS-1 method. Then, the Kimura distance between the sequences was calculated using the distmat function of the EMBOSS suite (*67*).

### Protein structure prediction and modeling

Structural models for OmpW proteins were generated using AlphaFold 3 (*68*). For each protein, the top-ranked model (labeled as “model 0”) was selected for downstream analysis based on its highest confidence score. Amino acid sequences were processed using SignalP (*CS*) and sequence comparison, to identify and remove the predicted signal peptides prior to structural modeling.

Structural alignments were performed using the US-align web server (*70*), which provides a sequence-independent alignment based on the TM-score and RMSD (Template Modeling score and Root-Mean-Square Deviation). Molecular visualization, surface characterization, and figure generation were conducted using PyMOL v3.1.6.1.

Internal channels and pore dimensions of the OmpW porins were characterized using the PoreWalker 1.0 server (*71*), based on the 3D models previously generated. To visualize the internal architecture and cavity distribution, cross-sectional views were generated at the XY-plane (bottom-left sectioning, considering only Z>0 coordinates). This analysis allowed for the identification of the pore centers and the assessment of the diameter profile along the channel axis. To map the pore diameter profile across the entire protein structure, transverse sections were taken at 1 Å intervals along the channel axis (sphere coordinates provided in the sphere2-fm.txt output file).

### Generation of *ompW* insertion mutants

To evaluate the functional roles of the *ompW1*, *ompW2a*, and *ompW3* paralogs, single-crossover insertion mutants were constructed in the *A. baumannii* ATCC 17978 reference strain. Briefly, internal fragments of each *ompW* protein-coding sequence were amplified by PCR using specific primers (Suppl. Table S3) and cloned into the pCR-Blunt II-TOPO vector (Thermo Fisher Scientific, Seville, Spain). Recombinant plasmids were propagated in competent *E. coli* DH5α cells (Invitrogen, Seville, Spain) via heat-shock transformation and subsequently purified using the QIAprep Spin Miniprep kit (Qiagen, Barcelona, Spain). The constructs were then introduced into electrocompetent *A. baumannii* ATCC 17978 cells by electroporation using a MicroPulser (Bio-Rad, Madrid, Spain). Following pulse delivery, bacteria were immediately recovered in warm SOC medium (Invitrogen), incubated under shaking conditions at 37°C for 1 hours, and finally plated on LB agar supplemented with 50 µg/mL kanamycin (Merck, Darmstadt, Germany) to select for stable transformants harboring the integrated plasmid.

### Construction of *ompW3/ompW2a* double mutant

To generate the double mutant *ompW3*::Amp *ompW2a*::Kan, the *ompW2a*::Kan insertion mutant was used as the parental strain. An additional plasmid was constructed from pCR™-Blunt II-TOPO®, which carried an internal fragment of ompW3, and an ampicillin resistance cassette was inserted using the restriction enzymes XcmI (New England Biolab, Barcelona, Spain). Selection was carried out on LB agar plates supplemented with 50 µg/mL ampicillin (Merck), allowing the isolation of colonies carrying both insertion cassettes. Integration of the second construct via single-crossover homologous recombination resulted in the simultaneous disruption of the *ompW2a* and *ompW3 loci*.

### Evaluation of mutant construction

Correct integration of the vector into the target genomic *locus* was verified by PCR (Suppl. Fig. S7). Definitive confirmation of the insertion sites was subsequently performed via Sanger sequencing (Suppl. Fig. S8), comparing the sequence chromatograms of the mutant strains against the wild-type reference genome.

### In vitro studies

All *in vitro* studies detailed below were carried out twice on different days to avoid bias.

### Individual and competitive growth curves

Growth curves of *A. baumannii* ATCC 17978 wild-type, *ompW1*::Kan, *ompW2a*::Kan, *ompW3*::Kan and *ompW3*::Amp *ompW2a*::Kan were determined by inoculating 20 mL of Luria Bertani (LB) broth with a starting concentration of 5,5 log_10_ CFU/mL. Concentration was then measured at different time points (2, 4, 8 and 24 hours) (*72*).

### Biofilm assay

Biofilm formation of *A. baumannii* ATCC 17978 wild-type, *ompW1*::Kan, *ompW2a*::Kan, *ompW3*::Kan and *ompW3*::Amp *ompW2a*::Kan was assessed inoculating 200 µL of each bacterial suspension at a concentration of 6 log_10_ CFU/mL into the wells of a 96-well tissue culture plate (Greiner Bio-One GmbH, Merck, Madrid, Spain) and incubated for 24 hours at 37°C to permit the formation of biofilm (*72*).

Subsequently, wells were washed three times with phosphate-buffered saline (PBS) to remove the cells that did not form biofilm and were not attached to the bottom of the well plates. Subsequently, biofilms were stained with 0.4% crystal violet (Merck) for 10 minutes at room temperature in the dark. Wash steps were performed again, to remove the residual crystal violet. Biofilm tinted by crystal violet was then solubilized with 200 µL of 96% ethanol for 15 minutes in the dark. Finally, biofilm biomass was quantified by measuring absorbance at 580 nm using a microplate reader (Asys UVM 340, Cambridge, UK).

### Surface motility test

Overnight cultures in LB of *A. baumannii* ATCC 17978 wild-type, *ompW1*::Kan, *ompW2a*::Kan, *ompW3*::Kan and *ompW3*::Amp *ompW2a*::Kan were adjusted to an optical density of 600 nm (OD600) in 1x PBS. Afterwards, 3 µL of the adjusted inoculum was plated in LB plates with 0,3% agarose (Sigma-Aldrich). Plates were incubated for 24 hours at 37°C, under 5% CO_2_ and a 90-95% relative humidity which promote bacterial surface motility (*72*).

### Bacterial adherence and invasion assays

For the bacterial adherence studies, human lung epithelial cells A549 (ATCC® CCL-185™) were seeded in 24 well plates at a concentration of 10^5^ cells/well for 24 hours, at 37°C. Overnight bacterial cultures of *A. baumannii* ATCC 17978 wild-type, *ompW1*::Kan, *ompW2a*::Kan, *ompW3*::Kan and *ompW3*::Amp *ompW2a*::Kan were diluted 1:1000 in PBS and added to the cells and incubated for 2 hours, at 37°C. Subsequently, 200 μl of trypsin (Gibco) were added and incubated for 5 minutes at 37°C. Afterwards, 200 μl of 0,5% Triton X-100 (Merck) were added and incubated for another 2 minutes, 37°C (*72*).

To determine bacterial invasion into A549 cells, these cells were incubated again with the same bacterial strains, under the conditions described above. After the incubation period, 256 μg/mL of amikacin (Sigma-Aldrich) were added to each well and the plates were further incubated for 30 min. Later, the plates was treated with trypsin and Triton X-100, as described previously (*72*).

### Animal use ethics

All animal procedures complied with the Guide for the Care and Use of Laboratory Animals and ARRIVE guidelines (https://arriveguidelines.org/). Experiments were approved by the Committee on the Ethics of Animal Experiments of the University Hospital of Virgen del Rocío, Seville, Spain, and from the Ministry of *Agricultura, Pesca y Desarrollo Rural* (07/02/2022/007). A murine model of peritoneal sepsis was established using immunocompetent female C57BL/6J mice of 7 weeks (20 g). Procedures were performed to minimize animal suffering, following the 3Rs principles.

### *In vivo* dissemination

Bacterial dissemination was analyzed for *A. baumannii* ATCC 17978 and the four isogenic *ompW* mutants (*ompW1*::Kan, *ompW2a*::Kan, *ompW3*::Kan and *ompW3*::Amp *ompW2a*::Kan). To this end, for each strain, mice were randomly selected and inoculated intraperitoneally with 0.5 ml of inoculum from each strain at a concentration equivalent to the MIC of the *A. baumannii* ATCC 17978 wild-type strain (8.5 log_10_ CFU/mL). Four mice of each group were randomly selected and sacrificed (sodium thiopental i.p.) 8 and 24 hours post-inoculation. Immediately after the sacrifice, different organs (spleen, lungs, kidneys, liver) and peritoneal fluid and blood, to quantify the bacterial load (log_10_ CFU/g and log_10_ CFU/mL) (*73*).

## Statistics

Statistical analyses were performed using R (version 4.3.2). The normal distribution of quantitative datasets was systematically evaluated using the Shapiro-Wilk test. For the *in vitro* phenotypic characterizations, as well as for the *in vivo* organ bacterial burden comparisons, statistical significance between the mutant strains and the wild-type control was determined using the non-parametric Mann-Whitney U test. To evaluate protein length variations between single-copy and multi-copy species, significant differences were determined using the Wilcoxon rank-sum test. For the molecular phylogeny divergence analyses, Pearson correlation coefficients (*r*) and their corresponding *p*-values were computed to assess the relationship between sequence distances and species divergence times. For multiple pairwise comparisons, *p*-values were formally adjusted using the Bonferroni correction method. A *p*-value lower than 0.05 was considered statistically significant (\**p* < 0.05, \*\**p* < 0.005, \*\*\**p* < 0.001).

## Acknowledgements

We thank C3UPO and CICA for the HPC support, and Zaida Pérez-Cano for the selection of bacterial species with multiple paralogs. Figures 5D and 6 were created with BioRender.com and Figure 8 contain elements created with it.

## Data availability

The code and input data used in this work are available at https://doi.org/10.5281/zenodo.21567000

## Supplementary Tables

**Suppl. Table S1.** Reference genomes of bacteria from the NCBI Genome with date March 2025. For each species, its genus, order, and family are listed, along with whether it was classified as Gram-negative or Gram-positive. The number of ompW genes found in its genome is also shown, as well as whether or not the species was found in the TimeTree database.

**Suppl. Table S2.** Chronological list of published studies investigating OmpW proteins in *A. baumannii*. For each study, the observed or hypothesized phenotypic roles are summarized. *Loci* are classified and relabeled as *ompW*, *ompW2*, or *ompW3* based on a retrospective analysis of the provided genomic data, or designated as ‘undetermined’ when the specific gene could not be verified.

**Suppl. Table S3.**
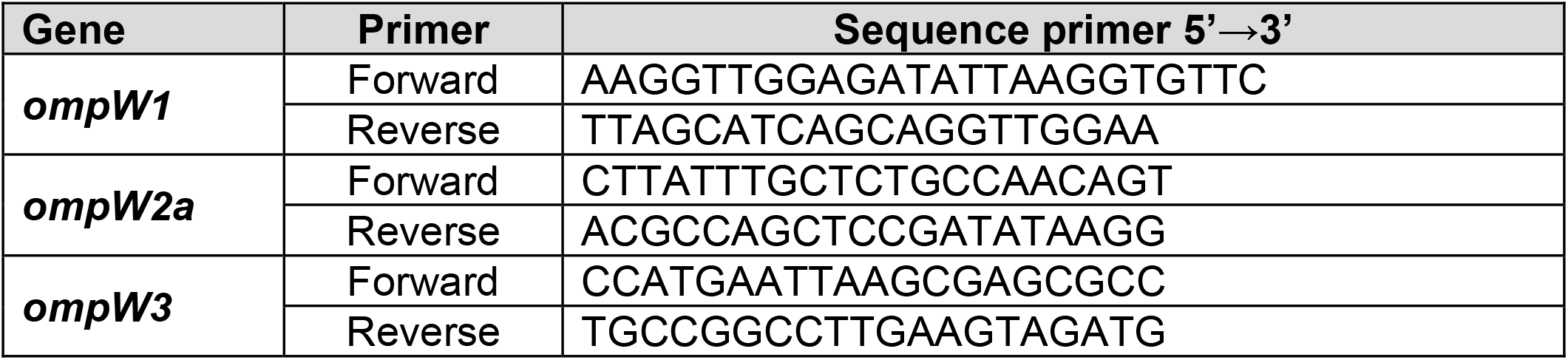
Primers used to amplify a fragment of the CDS of *ompW1*, *ompW2a* and *ompW3*, respectively to introduce it in pCRTM-Blunt II-TOPO vector.

## Supplementary Figures

**Suppl. Fig. S1.** Detailed phylogenetic distribution and conservation of six OmpW proteins within the *Burkholderiaceae*. The chronogram shows the evolutionary relationships among the analyzed genera of the *Burkholderiaceae*. The first heatmap shows the mean amino acid sequence identity of reciprocal best-hit orthologs to each of the six OmpW proteins from *Burkholderia stabilis* (A–F), averaged across the species analyzed within each genus. Gray cells indicate that no qualifying reciprocal ortholog was detected. The second heatmap shows the proportion of species within each genus containing a reciprocal ortholog of each OmpW protein. Numbers inside the cells indicate the number of species containing the gene relative to the total number of species analyzed for that genus, and color intensity represents the corresponding proportion. Labels A–F correspond to the OmpW groups defined in Fig. 2F. This family-level representation provides a detailed view of the broader phylogenetic distribution shown in Suppl. Fig. S2.

**Suppl. Fig. S2.** Phylogenetic distribution and conservation of six OmpW proteins across bacterial genera. The chronogram shows the evolutionary relationships among the analyzed genera, using one representative species per genus from the species-level TimeTree. Branch lengths represent time before present in millions of years (Mya). The first heatmap shows the mean amino acid sequence identity of reciprocal best-hit orthologs to each of the six OmpW proteins from *Burkholderia stabilis* (A–F), averaged across the species analyzed within each genus. Gray cells indicate that no qualifying reciprocal ortholog was detected. The second heatmap shows the proportion of species within each genus containing a reciprocal ortholog of each OmpW protein. Numbers inside the cells indicate the number of species containing the gene relative to the total number of species analyzed for that genus, and color intensity represents the corresponding proportion. Labels A-F correspond to the OmpW groups defined in Fig. 2F.

**Suppl. Fig. S3.** Prediction and genomic organization of the putative integrative and conjugative element carrying the horizontally acquired *ompW* variant in *Ralstonia mannitolilytica*. ICEfinder 2.0, implemented in the ICEberg 3.0 web server, predicted an attachment-site-delimited 116,760-bp mobile element spanning positions 1,596,574-1,713,333. The automated analysis classified the region as an integrative and mobilizable element (IME), as it detected a MOBH-family relaxase but did not identify a complete mating-pair formation system. Subsequent BLAST-based annotation identified additional homologs of Vir and conjugation-associated proteins, supporting its provisional classification as a putative integrative and conjugative element (ICE). The horizontally acquired B *ompW* copy is embedded within a naphthalene-degradation gene cluster. Functional categories are indicated by colors.

**Suppl. Fig. S4.** Sequence conservation of the genomic region encompassing the ompW2 gene *locus* in *A. baumannii*, including its flanking genes. The alignment incorporates the sequences corresponding to the four most common *ompW2a* and *ompW2b* variants. The same as Fig. 4C but including the sequence detail.

**Suppl. Fig. S5.** Multiple sequence alignment of *A. baumannii* OmpW paralogs (including the two highly divergent OmpW2 copies) and the *F. lacunae* OmpW ortholog. The primary sequence alignment was manually refined and guided by the 3D structural models of the six proteins to ensure accurate topological matching.

**Suppl. Fig. S6.** Sequence logo of the OmpW domain. The sequence logo illustrates the amino acid conservation profile across all sequences of the OmpW family available in the InterPro database (IPR005618). The visualization was generated using the Logomaker package.

**Suppl. Fig. S7.** Electroporated *A. baumannii* ATCC 17978 recovered in kanamicin/ampicillin plates (50 µg/mL). (A) GeneRuler 1 kb DNA Ladder (Thermo Fisher). (B) *ompW1*::Kan (998 bp). (C) *ompW2a*::Kan (1200 bp). (D) *ompW3*::Kan (765 bp). (E). *ompW3*::Amp ompW2a::Kan (765 bp). Primers used were Forward (Fw) M13 (align in the plasmid) and the corresponding Fw upstream for *ompW1*::Kan and *ompW2a*::Kan; and Fw intern for *ompW3*::Kan and *ompW3*::Amp *ompW2a*::Kan.

**Suppl. Fig. S8.** Sanger sequencing and schematic validation of cassette insertion sites in *ompW* disruption strains. For each targeted locus, the full coding sequence (CDS) is presented, with the region downstream of the insertion site highlighted in red to denote the frame disruption. Below each sequence, a linear genomic diagram illustrates the precise architectural impact of the integration: the unaffected upstream region (blue), the exact insertion site (orange arrow), and the downstream region affected by the insertion (grey). Junction sequences detailing the nucleotide boundaries at the insertion site are provided underneath each diagram.

## References

1. H. Hong, D. R. Patel, L. K. Tamm, B. van den Berg, The Outer Membrane Protein OmpW Forms an Eight-stranded β-Barrel with a Hydrophobic Channel. Journal of Biological Chemistry 281, 7568–7577 (2006).

2. X. Fu, J. Zhang, T. Li, M. Zhang, J. Li, B. Kan, The Outer Membrane Protein OmpW Enhanced V. cholerae Growth in Hypersaline Conditions by Transporting Carnitine. Front. Microbiol. 8 (2018).

3. M. Xiao, Y. Lai, J. Sun, G. Chen, A. Yan, Transcriptional Regulation of the Outer Membrane Porin Gene ompW Reveals its Physiological Role during the Transition from the Aerobic to the Anaerobic Lifestyle of Escherichia coli. Front. Microbiol. 7 (2016).

4. C. Seddon, G. Frankel, K. Beis, Structure of the outer membrane porin OmpW from the pervasive pathogen Klebsiella pneumoniae. Acta Cryst F 80, 22–27 (2024).

5. X.-B. Wu, L.-H. Tian, H.-J. Zou, C.-Y. Wang, Z.-Q. Yu, C.-H. Tang, F.-K. Zhao, J.-Y. Pan, Outer membrane protein OmpW of Escherichia coli is required for resistance to phagocytosis. Research in Microbiology 164, 848–855 (2013).

6. W. W. Low, C. Seddon, K. Beis, G. Frankel, The Interaction of the F-Like Plasmid-Encoded TraN Isoforms with Their Cognate Outer Membrane Receptors. Journal of Bacteriology 205, e00061–23 (2023).

7. B. Liu, J. Wang, J. Hu, L. Li, L. Liu, OmpW overexpression in meropenem-exposed Acinetobacter baumannii persister cells enhances virulence and reveals a candidate target. Front. Microbiol. 16 (2025).

8. M. Catel-Ferreira, S. Marti, L. Guillon, L. Jara, G. Coadou, V. Molle, E. Bouffartigues, G. Bou, I. Shalk, T. Jouenne, X. Vila-Farrés, E. Dé, The outer membrane porin OmpW of Acinetobacter baumannii is involved in iron uptake and colistin binding. FEBS Letters 590, 224–231 (2016).

9. H. Pilsl, D. Smajs, V. Braun, Characterization of Colicin S4 and Its Receptor, OmpW, a Minor Protein of the Escherichia coli Outer Membrane. Journal of Bacteriology 181, 3578–3581 (1999).

10. D. Xu, J. Zhang, J. Liu, J. Xu, H. Zhou, L. Zhang, J. Zhu, B. Kan, Outer Membrane Protein OmpW Is the Receptor for Typing Phage VP5 in the Vibrio cholerae O1 El Tor Biotype. Journal of Virology 88, 7109–7111 (2014).

11. C. Zhu, S. Liang, N. Yang, S. Li, J. Xue, R. Zhou, X. Hong, S. Chen, N. Gao, Q. Du, J. Huang, G. Feng, X. Du, Immunogenicity-Guided Design of an Acinetobacter baumanii Vaccine. European Journal of Immunology 55, e70019 (2025).

12. S. McClean, M. E. Healy, C. Collins, S. Carberry, L. O’Shaughnessy, R. Dennehy, Á. Adams, H. Kennelly, J. M. Corbett, F. Carty, L. A. Cahill, M. Callaghan, K. English, B. P. Mahon, S. Doyle, M. Shinoy, Linocin and OmpW Are Involved in Attachment of the Cystic Fibrosis-Associated Pathogen Burkholderia cepacia Complex to Lung Epithelial Cells and Protect Mice against Infection. Infect Immun 84, 1424–1437 (2016).

13. I. Soojhawon, N. Pattabiraman, A. Tsang, A. L. Roth, E. Kang, S. M. Noble, Discovery of novel inhibitors of multidrug-resistant *Acinetobacter baumannii*. Bioorganic & Medicinal Chemistry 25, 5477–5482 (2017).

14. S. Shahryari, P. Mohammadnejad, K. A. Noghabi, Screening of anti-Acinetobacter baumannii phytochemicals, based on the potential inhibitory effect on OmpA and OmpW functions. R Soc Open Sci. 8, 201652 (2021).

15. A. Y. Peleg, H. Seifert, D. L. Paterson, Acinetobacter baumannii: Emergence of a Successful Pathogen. Clin Microbiol Rev 21, 538–582 (2008).

16. E. Tacconelli, E. Carrara, A. Savoldi, S. Harbarth, M. Mendelson, D. L. Monnet, C. Pulcini, G. Kahlmeter, J. Kluytmans, Y. Carmeli, M. Ouellette, K. Outterson, J. Patel, M. Cavaleri, E. M. Cox, C. R. Houchens, M. L. Grayson, P. Hansen, N. Singh, U. Theuretzbacher, N. Magrini, WHO Pathogens Priority List Working Group, Discovery, research, and development of new antibiotics: the WHO priority list of antibiotic-resistant bacteria and tuberculosis. Lancet Infect Dis 18, 318–327 (2018).

17. A. Siroy, P. Cosette, D. Seyer, C. Lemaître-Guillier, D. Vallenet, A. Van Dorsselaer, S. Boyer-Mariotte, T. Jouenne, E. Dé, Global comparison of the membrane subproteomes between a multidrug-resistant Acinetobacter baumannii strain and a reference strain. J Proteome Res 5, 3385–3398 (2006).

18. S.-H. Yun, C.-W. Choi, S.-H. Park, J. C. Lee, S.-H. Leem, J.-S. Choi, S. Kim, S. I. Kim, Proteomic analysis of outer membrane proteins from Acinetobacter baumannii DU202 in tetracycline stress condition. J Microbiol. 46, 720–727 (2008).

19. J. Provasi Cardoso, R. Cayô, R. Girardello, A. C. Gales, Diversity of mechanisms conferring resistance to β-lactams among OXA-23–producing *Acinetobacter baumannii* clones. Diagnostic Microbiology and Infectious Disease 85, 90–97 (2016).

20. M. L. Gil-Marqués, J. Pachón, Y. Smani, iTRAQ-Based Quantitative Proteomic Analysis of Acinetobacter baumannii under Hypoxia and Normoxia Reveals the Role of OmpW as a Virulence Factor. Microbiology Spectrum 10, e02328–21 (2022).

21. B. L. Schmitt, B. F. Leal, M. Leyser, M. P. de Barros, D. S. Trentin, C. A. S. Ferreira, S. D. de Oliveira, Increased ompW and ompA expression and higher virulence of Acinetobacter baumannii persister cells. BMC Microbiol 23, 157 (2023).

22. J. Park, M. Kim, B. Shin, M. Kang, J. Yang, T. K. Lee, W. Park, A novel decoy strategy for polymyxin resistance in Acinetobacter baumannii. Elife 10, e66988 (2021).

23. J. Boral, C. Vatansever, G. Ozcan, S. Keske, S. Menekse, M. Gonen, F. Can, Resensitization to colistin results in rapid and stable recovery of adherence, serum resistance and ompW in Acinetobacter baumannii. PLoS One 19, e0309307 (2024).

24. E. V. Koonin, Orthologs, paralogs, and evolutionary genomics. Annu. Rev. Genet. 3G, 309–338 (2005).

25. J. Botelho, H. Schulenburg, The Role of Integrative and Conjugative Elements in Antibiotic Resistance Evolution. Trends Microbiol 29, 8–18 (2021).

26. J. M. Yagi, D. Sims, T. Brettin, D. Bruce, E. L. Madsen, The genome of Polaromonas naphthalenivorans strain CJ2, isolated from coal tar-contaminated sediment, reveals physiological and metabolic versatility and evolution through extensive horizontal gene transfer. Environmental Microbiology 11, 2253–2270 (2009).

27. J. Ma, Y. Zhai, Y. Cui, G. Gao, M. Ying, Y. Zhao, A. Antunes, L. Huang, M. Li, Study and Modification of the Polycyclic Aromatic Hydrocarbon Degradation Gene Cluster in Burkholderia sp. FM-2. Microorganisms 13, 2079 (2025).

28. T. J. Treangen, E. P. C. Rocha, Horizontal Transfer, Not Duplication, Drives the Expansion of Protein Families in Prokaryotes. PLOS Genetics 7, e1001284 (2011).

29. S. M. Soucy, J. Huang, J. P. Gogarten, Horizontal gene transfer: building the web of life. Nat Rev Genet 16, 472–482 (2015).

30. B. J. Arnold, I.-T. Huang, W. P. Hanage, Horizontal gene transfer and adaptive evolution in bacteria. Nat Rev Microbiol 20, 206–218 (2022).

31. W. Li, L. Wen, C. Li, R. Chen, Z. Ye, J. Zhao, J. Pan, Contribution of the outer membrane protein OmpW in Escherichia coli to complement resistance from binding to factor H. Microb Pathog 98, 57–62 (2016).

32. X. Chen, Z. Shao, L. Wu, B. He, W. Yang, J. Chen, E. Jin, Q. Huang, L. Lei, J. Xu, H. Li, H. Zhang, Y. Wan, W. Liu, R. Zhou, Involvement of the Actinobacillus pleuropneumoniae ompW Gene in Confrontation of Environmental Pressure. Front. Vet. Sci. **9** (2022).

33. X. Cao, C. Cao, Z. Chen, J. Li, Z. Yao, Y. Zheng, J. Wu, Z. Li, Y. Hu, G. Hao, G. Zhu, W. Köster, A. P. White, Y. Wang, Genome-wide investigation of outer membrane protein families under mosaic evolution in Escherichia coli. Applied and Environmental Microbiology 0, e00557–25 (2025).

34. X. Chen, X. Cai, Z. Chen, J. Wu, G. Hao, Q. Luo, S. Liu, J. Zhang, Y. Hu, G. Zhu, W. Koester, A. P. White, Y. Cai, Y. Wang, Mosaic Evolution of Beta-Barrel-Porin-Encoding Genes in Escherichia coli. Appl Environ Microbiol 88, e0006022 (2022).

35. S. Abdollahi, Z. Raoufi, A novel vaccine candidate against *A. baumannii* based on a new OmpW family protein (OmpW2); structural characterization, antigenicity and epitope investigation, and in-vivo analysis. Microbial Pathogenesis 183, 106317 (2023).

36. C. Rumbo, E. Gato, M. López, C. Ruiz de Alegría, F. Fernández-Cuenca, L. Martínez-Martínez, J. Vila, J. Pachón, J. M. Cisneros, J. Rodríguez-Baño, A. Pascual, G. Bou, M. Tomás, on behalf of the Spanish Group of Nosocomial Infections and Mechanisms of Action and Resistance to Antimicrobials (GEIH-GEMARA) from the Spanish Society of Clinical Microbiology and Infectious Diseases (SEIMC) and the Spanish Network for Research in Infectious Diseases (REIPI), Contribution of Efflux Pumps, Porins, and β-Lactamases to Multidrug Resistance in Clinical Isolates of Acinetobacter baumannii. Antimicrobial Agents and Chemotherapy 57, 5247–5257 (2013).

37. Z. Y. Kho, M. A. K. Azad, Y. Zhu, M.-L. Han, Q. (Tony) Zhou, T. Velkov, T. Naderer, J. Li, Transcriptomic interplay between Acinetobacter baumannii, human macrophage and polymyxin. bioRxiv [Preprint] (2024). 10.1101/2024.01.23.576770.

38. V. Tiwari, J. Vashistt, A. Kapil, R. R. Moganty, Comparative Proteomics of Inner Membrane Fraction from Carbapenem-Resistant Acinetobacter baumannii with a Reference Strain. PLOS ONE 7, e39451 (2012).

39. S. Chopra, K. Ramkissoon, D. C. Anderson, A systematic quantitative proteomic examination of multidrug resistance in *Acinetobacter baumannii*. Journal of Proteomics 84, 17–39 (2013).

40. S.-Y. Lee, M. H. Oh, S. H. Yun, C.-W. Choi, E. C. Park, H. S. Song, H. Lee, Y.-S. Yi, J. Shin, C. Chung, J. Y. Moon, J. C. Lee, G.-H. Kim, S. I. Kim, Genomic characterization of extensively drug-resistant Acinetobacter baumannii strain, KAB03 belonging to ST451 from Korea. Infection, Genetics and Evolution 65, 150– 158 (2018).

41. Y.-K. Tsai, C.-H. Liou, J.-C. Lin, C.-P. Fung, F.-Y. Chang, L. K. Siu, Effects of different resistance mechanisms on antimicrobial resistance in *Acinetobacter baumannii*: a strategic system for screening and activity testing of new antibiotics. International Journal of Antimicrobial Agents 55, 105918 (2020).

42. S. Kashyap, P. Sharma, N. Capalash, Tobramycin Stress Induced Differential Gene Expression in Acinetobacter baumannii. Curr Microbiol 7**G**, 88 (2022).

43. W. M. Nageeb, N. AlHarbi, A. A. Alrehaili, S. A. Zakai, A. Elfadadny, H. F. Hetta, Global genomic epidemiology of chromosomally mediated non-enzymatic carbapenem resistance in Acinetobacter baumannii: on the way to predict and modify resistance. Front. Microbiol. 14 (2023).

44. Y. Boulaamane, I. Molina Panadero, A. Hmadcha, C. Atalaya Rey, S. Baammi, A. El Allali, A. Maurady, Y. Smani, Antibiotic discovery with artificial intelligence for the treatment of Acinetobacter baumannii infections. mSystems **9**, e00325–24 (2024).

45. I. Molina-Panadero, M. Morales-Tenorio, A. García-Rubia, T. Ginex, K. Eskandari, A. Martinez, C. Gil, Y. Smani, Discovery of new antimicrobial thiophene derivatives with activity against drug-resistant Gram negative-bacteria. Front. Pharmacol. 15 (2024).

46. P. Pumirat, S. Santajit, W. Tunyong, T. Kong-Ngoen, S. Tandhavanant, S. Lohitthai, A. Rungruengkitkun, N. Chantratita, S. Ampawong, O. Reamtong, N. Indrawattana, Impact of AbaI mutation on virulence, biofilm development, and antibiotic susceptibility in Acinetobacter baumannii. Sci Rep 14, 21521 (2024).

47. X. Li, D. Feng, J. Zhou, W. Wu, C. He, W. Gan, W. Zheng, B. Peng, T. Zhang, Synergy of ATP and Meropenem in Stimulating the TCA Cycle to Enhance Killing of Carbapenem-Resistant Acinetobacter baumannii. Microbial Biotechnology 18, e70199 (2025).

48. S.-H. Yun, C.-W. Choi, S.-O. Kwon, G. W. Park, K. Cho, K.-H. Kwon, J. Y. Kim, J. S. Yoo, J. C. Lee, J.-S. Choi, S. Kim, S. I. Kim, Quantitative Proteomic Analysis of Cell Wall and Plasma Membrane Fractions from Multidrug-Resistant Acinetobacter baumannii. J. Proteome Res. 10, 459–469 (2011).

49. S. P. Bhamidimarri, M. Zahn, J. D. Prajapati, C. Schleberger, S. Söderholm, J. Hoover, J. West, U. Kleinekathöfer, D. Bumann, M. Winterhalter, B. van den Berg, A Multidisciplinary Approach toward Identification of Antibiotic Scaffolds for Acinetobacter baumannii. Structure 27, 268–280.e6 (2019).

50. C. C. Nwugo, J. A. Gaddy, D. L. Zimbler, L. A. Actis, Deciphering the iron response in *Acinetobacter baumannii*: A proteomics approach. Journal of Proteomics 74, 44–58 (2011).

51. D. Fu, Q. Zheng, X. Wu, J. Wu, Y. Shao, Z. Wang, J. Tu, X. Song, K. Qi, The transcriptional regulator EtrA mediates ompW contributing to the pathogenicity of avian pathogenic Escherichia coli. Vet Microbiol 283, 109775 (2023).

52. D. Scribano, E. Cheri, A. Pompilio, G. Di Bonaventura, M. Belli, M. Cristina, L. Sansone, C. Zagaglia, M. Sarshar, A. T. Palamara, C. Ambrosi, Acinetobacter baumannii OmpA-like porins: functional characterization of bacterial physiology, antibiotic-resistance, and virulence. Commun Biol 7, 948 (2024).

53. P. Rassam, N. A. Copeland, O. Birkholz, C. Tóth, M. Chavent, A. L. Duncan, S. J. Cross, N. G. Housden, R. Kaminska, U. Seger, D. M. Quinn, T. J. Garrod, M. S. P. Sansom, J. Piehler, C. G. Baumann, C. Kleanthous, Supramolecular assemblies underpin turnover of outer membrane proteins in bacteria. Nature 523, 333–336 (2015).

54. K. M. Clemmer, R. A. Bonomo, P. N. Rather, Genetic analysis of surface motility in Acinetobacter baumannii. Microbiology 157, 2534–2544 (2011).

55. I. Schober, J. Koblitz, J. Sardà Carbasse, C. Ebeling, M. L. Schmidt, A. Podstawka, R. Gupta, V. Ilangovan, J. Chamanara, J. Overmann, L. C. Reimer, BacDive in 2025: the core database for prokaryotic strain data. Nucleic Acids Res 53, D748–D756 (2025).

56. S. Kumar, M. Suleski, J. M. Craig, A. E. Kasprowicz, M. Sanderford, M. Li, G. Stecher, S. B. Hedges, TimeTree 5: An Expanded Resource for Species Divergence Times. Mol Biol Evol 3**G**, msac174 (2022).

57. O. Schwengers, L. Jelonek, M. A. Dieckmann, S. Beyvers, J. Blom, A. Goesmann, Bakta: rapid and standardized annotation of bacterial genomes via alignment-free sequence identification. Microbial Genomics 7, 000685 (2021).

58. R. D. Finn, J. Clements, S. R. Eddy, HMMER web server: interactive sequence similarity searching. Nucleic Acids Res, doi: 10.1093/nar/gkr367 (2011).

59. M. Blum, H.-Y. Chang, S. Chuguransky, T. Grego, S. Kandasaamy, A. Mitchell, G. Nuka, T. Paysan-Lafosse, M. Qureshi, S. Raj, L. Richardson, G. A. Salazar, L. Williams, P. Bork, A. Bridge, J. Gough, D. H. Haft, I. Letunic, A. Marchler-Bauer, H. Mi, D. A. Natale, M. Necci, C. A. Orengo, A. P. Pandurangan, C. Rivoire, C. J. A. Sigrist, I. Sillitoe, N. Thanki, P. D. Thomas, S. C. E. Tosatto, C. H. Wu, A. Bateman, R. D. Finn, The InterPro protein families and domains database: 20 years on. Nucleic Acids Research 4**G**, D344–D354 (2021).

60. K. Katoh, D. M. Standley, MAFFT multiple sequence alignment software version 7: improvements in performance and usability. Mol. Biol. Evol. 30, 772–780 (2013).

61. J. L. Steenwyk, T. J. B. Iii, Y. Li, X.-X. Shen, A. Rokas, ClipKIT: A multiple sequence alignment trimming software for accurate phylogenomic inference. PLOS Biology 18, e3001007 (2020).

62. B. Q. Minh, H. A. Schmidt, O. Chernomor, D. Schrempf, M. D. Woodhams, A. von Haeseler, R. Lanfear, IQ-TREE 2: New Models and Efficient Methods for Phylogenetic Inference in the Genomic Era. Mol Biol Evol 37, 1530–1534 (2020).

63. M. Wang, G. Liu, M. Liu, C. Tai, Z. Deng, J. Song, H.-Y. Ou, ICEberg 3.0: functional categorization and analysis of the integrative and conjugative elements in bacteria. Nucleic Acids Res 52, D732–D737 (2024).

64. A. Moreno-Rodríguez, A. Rubio, A. Garzón, Y. Smani, A. J. Pérez-Pulido, The evolutionary replacement of restriction-modification by Ssp antiviral systems is associated with the distribution of prophages in the major clonal group of Acinetobacter baumannii. mBio 0, e02135–25 (2025).

65. L. Fu, B. Niu, Z. Zhu, S. Wu, W. Li, CD-HIT: accelerated for clustering the next-generation sequencing data. Bioinformatics 28, 3150–3152 (2012).

66. S. Xu, L. Li, X. Luo, M. Chen, W. Tang, L. Zhan, Z. Dai, T. T. Lam, Y. Guan, G. Yu, Ggtree: A serialized data object for visualization of a phylogenetic tree and annotation data. iMeta 1, e56 (2022).

67. P. Rice, I. Longden, A. Bleasby, EMBOSS: the European Molecular Biology Open Software Suite. Trends Genet 16, 276–277 (2000).

68. J. Abramson, J. Adler, J. Dunger, R. Evans, T. Green, A. Pritzel, O. Ronneberger, L. Willmore, A. J. Ballard, J. Bambrick, S. W. Bodenstein, D. A. Evans, C.-C. Hung, M. O’Neill, D. Reiman, K. Tunyasuvunakool, Z. Wu, A. Žemgulytė, E. Arvaniti, C. Beattie, O. Bertolli, A. Bridgland, A. Cherepanov, M. Congreve, A. I. Cowen-Rivers, A. Cowie, M. Figurnov, F. B. Fuchs, H. Gladman, R. Jain, Y. A. Khan, C. M. R. Low, K. Perlin, A. Potapenko, P. Savy, S. Singh, A. Stecula, A. Thillaisundaram, C. Tong, S. Yakneen, E. D. Zhong, M. Zielinski, A. Žídek, V. Bapst, P. Kohli, M. Jaderberg, D. Hassabis, J. M. Jumper, Accurate structure prediction of biomolecular interactions with AlphaFold 3. Nature 630, 493–500 (2024).

69. H. Nielsen, F. Teufel, S. Brunak, G. von Heijne, SignalP: The Evolution of a Web Server. Methods Mol Biol 2836, 331–367 (2024).

70. C. Zhang, M. Shine, A. M. Pyle, Y. Zhang, US-align: universal structure alignments of proteins, nucleic acids, and macromolecular complexes. Nat Methods 1**G**, 1109–1115 (2022).

71. M. Pellegrini-Calace, T. Maiwald, J. M. Thornton, PoreWalker: A Novel Tool for the Identification and Characterization of Channels in Transmembrane Proteins from Their Three-Dimensional Structure. PLOS Computational Biology 5, e1000440 (2009).

72. S. Herrera-Espejo, J. L. Domínguez-Miranda, J. I. Rodríguez-Mogollo, J. Pachón, E. Cordero, M. E. Pachón-Ibáñez, Effects of pH on the Pathogenicity of Escherichia coli and Klebsiella pneumoniae on the Kidney: In Vitro and In Vivo Studies. International Journal of Molecular Sciences 25, 7925 (2024).

73. G. Labrador-Herrera, A. J. Pérez-Pulido, R. Álvarez-Marín, C. S. Casimiro-Soriguer, T. Cebrero-Cangueiro, J. Morán-Barrio, J. Pachón, A. M. Viale, M. E. Pachón-Ibáñez, Virulence role of the outer membrane protein CarO in carbapenem-resistant Acinetobacter baumannii. Virulence 11, 1727–1737 (2020).

